# PlasRAG: comprehensive plasmid characterization and retrieval through sequence-text alignment

**DOI:** 10.1101/2025.06.22.660968

**Authors:** Yongxin Ji, Herui Liao, Jiaojiao Guan, Jiayu Shang, Yanni Sun

## Abstract

Plasmids play a pivotal role in the emergence of multidrug-resistant and pathogenic bacteria, posing significant clinical challenges. The integration of metagenomic sequencing with advanced bioinformatics tools surpasses traditional wet lab methods, leading to the discovery of millions of plasmids from diverse origins. However, the rapidly growing number of unannotated plasmids necessitates comprehensive characterization of their multi-faceted properties, such as risk indices and ecological contexts, to support various downstream applications. Achieving this goal is hindered by several challenges, including the limited availability of plasmid characterization tools, the inadequacies of alignment-based methods for novel plasmids, and inconsistencies in manual annotations across plasmid reference databases. To address these issues, we present PlasRAG, a novel tool that integrates two key modules: multi-faceted property characterization of query plasmids and plasmid DNA retrieval based on textual queries. At its core, PlasRAG employs a bidirectional multi-modal information retrieval model that aligns DNA sequences with textual data, effectively overcoming the limitations of traditional approaches. Specifically, within the characterization module, PlasRAG leverages the retrieval-augmented generation (RAG) framework and the Llama-3 large language model (LLM) to provide accurate and context-aware responses to user queries. Rigorous experiments demonstrate that PlasRAG delivers robust performance and enhanced analytical capabilities, underscoring the effectiveness of its architectural design. In particular, experiments on a real-world plasmid dataset curated from diverse human gut metagenomes suggest that plasmids with a broader host range and encoded ARGs tend to spread more extensively. The source code of PlasRAG is available via: https://github.com/Orin-beep/PlasRAG.

## Introduction

A plasmid is typically a small, circular replicon found in bacteria. Unlike large chromosomes that encode essential genetic information for bacterial survival, plasmids carry additional genes that confer accessory traits to their hosts, offering selective advantages in various environmental conditions. For example, plasmids that carry antibiotic resistance genes (ARGs) can protect their hosts from antibiotics, and those containing virulence factor genes (VFGs) can contribute to bacterial pathogenicity. Moreover, as a type of mobile genetic element (MGE), plasmids can mediate horizontal gene transfer by transferring between different bacterial cells through a process known as conjugation [1]. Consequently, plasmid dissemination within bacterial populations is a primary driver of the emergence and rapid evolution of multidrug-resistant clinical pathogens [2], which in turn leads to bacterial disease outbreaks [3]. Given their significant clinical implications, extensive efforts have been made to investigate plasmids, focusing on their biological functions and ecology, particularly how they interact with other entities (such as bacteria and MGEs) within the same community [4].

Initially, researchers employed culture-dependent methods to obtain plasmids, isolating them from cultured bacteria based on their molecular characteristics [5]. However, in addition to being labor-intensive and time-consuming, this wet lab approach also results in a low diversity of isolated plasmids, underrepresenting those derived from non-culturable and slow-growing bacteria. With the advancement of sequencing technologies, integrating metagenomic sequencing with computational plasmid identification tools offers a more robust pipeline than traditional methods. First, the rapidly increasing metagenomic data catalogues a wide microbial diversity that serves as a valuable source for identifying plasmids. For instance, the IMG/M dataset [6] has curated 38,503 metagenomes from 37 subcategories of ecosystems, including wastewater, air, and human-associated ecosystems. Second, several methods, such as geNomad [7] and PLASMe [8], have been developed to identify plasmids from metagenomes effectively. Typically, these tools utilize deep learning (DL) models to distinguish plasmid DNA from other entities by using marker proteins or sequence motifs as features. Building on available plasmid identification methods, two high-quality plasmid databases, IMG/PR [9] and PIPdb [10], have collectively curated nearly 1.5 million novel plasmids. The large-scale plasmid data provide a solid foundation for in-depth plasmid analysis.

Given the rapid accumulation of newly discovered plasmids, there is an urgent need to characterize their properties from multiple facets, thereby facilitating downstream research. We highlight the importance of this process from four perspectives. First, the host-beneficial traits carried by plasmids, such as antimicrobial resistance (AMR), virulence factors (VFs), and heavy metal resistance (HMR), are major contributors to multidrug-resistant (MDR) infectious diseases. Plasmids with these traits are considered high-risk, typically posing significant threats to public health. By identifying high-risk plasmids, various treatment strategies, such as degrading targeted plasmids [11], can effectively control related clinical outbreaks of bacterial infections. Second, examining basic plasmid properties, including replication, stability, mobility, and host range, can contribute to tracking their transmission pathways within bacterial populations, which drive the rapid evolution of bacteria. Third, predicting the ecosystems associated with plasmids is significant because plasmids serve as the primary vehicles that enhance bacterial adaptation to changing environmental conditions. In summary, a computational tool that comprehensively characterizes input plasmid DNA should be developed.

There are three challenges to achieving comprehensive plasmid characterization. First, it is beneficial to employ more available high-quality plasmid databases, such as IMG/PR, PIPdb, and PLSDB [12], which can provide a strong data foundation for designing new methods. Nevertheless, directly merging them into a unified database is challenging, as they contain raw property annotations from diverse perspectives and descriptive styles (crowd-sourced labeling). For example, while IMG/PR annotates plasmids sourced from urban environments, the other two databases lack this information in their annotated vocabularies. Second, only a few tools have been proposed to characterize plasmids from limited facets, such as MOSTPLAS [13] for predicting host range, MOBFinder [14] for classifying mobilization types, and PlasmidFinder [15] for identifying replicons. All of these tools were developed using NCBI plasmids, which have limited data size and diversity. Additionally, since different property facets are associated with various plasmid sequence features and can be formulated as different problem types (e.g., classification or regression), we need a versatile framework to integrate the multi-faceted plasmid characterization tasks. Third, while alignment-based methods like BLAST can be adapted to annotate a query plasmid by identifying the most similar reference plasmid in the database, they struggle with novel plasmids that lack reliable alignments. Moreover, we cannot guarantee that two similar plasmids share all properties.

In addition to characterizing plasmid properties, retrieving eligible plasmid sequences is also valuable. By selecting retrieval queries from a comprehensive property vocabulary, users can accurately identify the plasmids that match their requirements from a given candidate set, thereby facilitating the plasmid-related sequencing data analysis. PlasmidScope [16] is an integrated plasmid database constructed from ten existing plasmid databases and features a plasmid sequence retrieval function. However, PlasmidScope has limitations in coarse search conditions, such as specifying bacterial hosts at the phylum level or determining whether plasmids encode ARGs or VFs. Additionally, it can only return plasmids from its pre-indexed database and cannot handle user-provided query plasmids. Therefore, in this work, we introduce a tool called PlasRAG, which serves two purposes: (1) plasmid property characterization, and (2) plasmid DNA retrieval. Both purposes are based on the same bidirectional information retrieval (IR) model, which can predict properties for input plasmids (forward) and search for matched plasmids based on query properties (reverse). We expand the scope of plasmid characterization by incorporating the retrieval-augmented generation (RAG) framework [17]. Specifically, when provided with a query plasmid DNA sequence, we utilize the properties and the most relevant literature predicted by the IR model as external knowledge. By integrating this retrieved knowledge with the user’s textual query (e.g., a question about the query plasmid), we leverage the text summarization and reasoning capabilities of a powerful large language model (LLM) to generate more accurate and reliable responses. As illustrated in Fig. 1, this approach transforms plasmid characterization into an interactive human–machine framework, enabling more effective a nd u ser-friendly plasmid analysis.

**Fig. 1.**
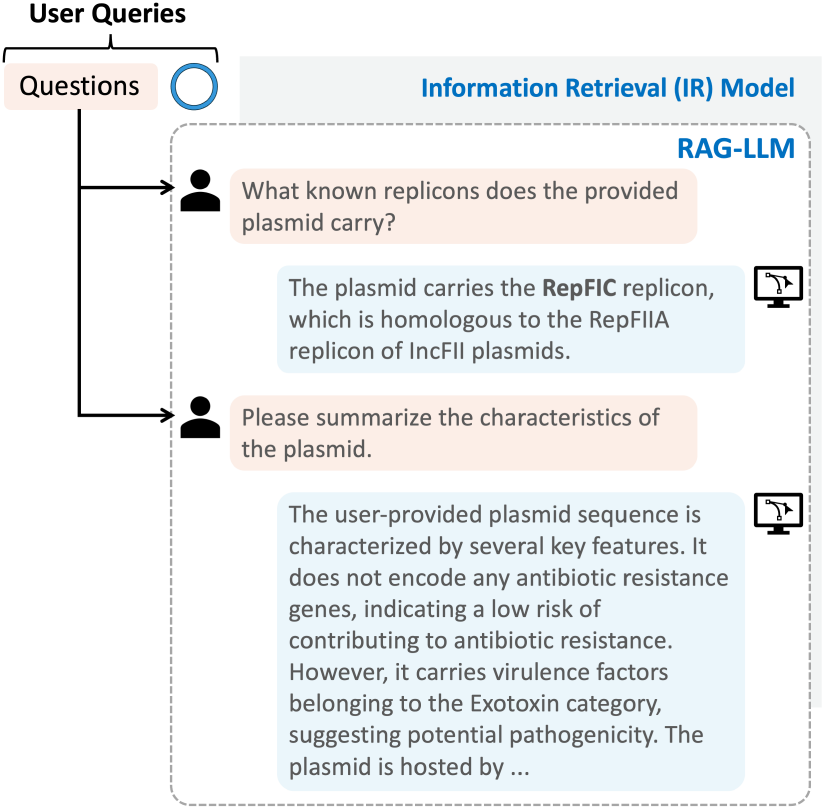
A simplified e xample of the PlasRAG’sp lasmid characterization purpose. Users can provide two queries: one plasmid DNA sequence (represented by the blue circle) and related questions (labeled with an orange background). PlasRAG will generate reliable responses using the retrieval-augmented generation (RAG) framework, which utilizes the information retrieval (IR) model to search for ‘external knowledge’ about the plasmid.

### Overview of PlasRAG

Our proposed solution to address the above challenges involves extracting semantic features from the textual properties, rather than treating each property as a classification label. The textual features can effectively bridge the annotation inconsistency between different plasmid databases, as they reflect semantic similarities between annotations. With this approach, we can implement the bidirectional IR model based on a multi-modal learning algorithm that learns feature associations between plasmid DNA in biological sequence modality and their properties in textual modality. Within the IR model, a sequence encoder and a text encoder serve as two foundational components that extract meaningful representations for plasmid DNA and textual properties, respectively. The multi-modal architecture offers three key advantages. First, instead of using generic language models (LMs) from the natural language processing (NLP) field, we prefer domain-specific biomedical LMs, as they effectively capture specialized relationships between plasmid properties, thus providing additional informative features. Additionally, our proposed framework is versatile and can be easily adapted to various problem types of property facets without requiring specific modifications. Second, a well-trained sequence encoder is more flexible in characterizing plasmid DNA, particularly those with novel sequence compositions, which addresses the limitations of alignment-based methods like BLAST. Third, the RAG framework effectively minimizes response hallucination. This is achieved by allowing the IR model and LLM to perform their respective functions: the IR model captures DNA-text associations, while the LLM focuses exclusively on text summarization and question answering.

### Related work

Several efforts have been made to explore multi-modal learning algorithms that connect features from biological sequences (DNA or proteins) and texts. Among them, protein-text pattern learning is a promising research area, as the high-quality Swiss-Prot database [18] contains extensive annotations in natural language for protein sequences. ProtST [19] is a representative work that aims to enhance protein understanding by integrating biomedical texts describing protein functions and other properties. Through implementing a cross-attention-based fusion model that learns associations between amino acid and word representations, the pre-trained ProtST can be adapted for various downstream tasks, such as fitness landscape prediction and functional protein retrieval. However, the ProtST method cannot be directly generalized to plasmids, as DNA is much longer than proteins, making DNA modeling a significant challenge. One common trait of ProtST and our proposed PlasRAG is that both utilize two types of encoder language models (LMs), namely LLMs for encoding natural language and biological sequence LMs, as key components of their methods.

### LLMs in biomedical domain

Several pre-trained LLMs have been developed to optimize NLP tasks in the biomedical domain by extracting specialized features from biomedical texts. As the first Transformer-based [20] biomedical LM, BioBERT [21] was initialized from the original BERT [22] and further pre-trained on two life science literature databases: PubMed abstracts [23] and full-text articles from PMC [24], encompassing a total of 18 billion words. The domain-specific training allows BioBERT to outperform the original BERT in various biomedical text-mining tasks. However, Gu et al. proposed that the general texts outside the biomedical domain, which contributed to the training of the original BERT, negatively impact the performance of BioBERT. Thus, they developed the BERT-based BioMedBERT model [25] entirely from scratch, constructing its vocabulary and conducting pre-training exclusively on PubMed abstracts. Their experiments validated that BioMedBERT demonstrates a strong ability to understand biomedical terms, achieving new state-of-the-art (SOTA) performance compared to other models, including BioBERT. Many other works have also been introduced based on BioMedBERT, adapting advanced training methodologies to address various research problems. For example, Jin et al. designed a biomedical IR tool called MedCPT [26], which can retrieve relevant literature based on user-input query sentences. Its central algorithm employs contrastive learning on query-article user interaction data from PubMed. MedCPT features two encoders, both initialized from BioMedBERT: one encodes user queries, while the other encodes article abstracts. After training on 255 million PubMed query-article search logs, MedCPT demonstrates reliable performance in literature retrieval for zero-shot user queries.

### Biological sequence LMs

Biological sequence LMs can be categorized into protein language models (PLMs) and DNA foundation models. Both of them formulate biological sequences into languages of life defined by their respective fundamental units. For example, PLMs typically treat each amino acid as a token and consider the entire protein as a sentence. After pretraining on large-scale protein sequence corpora (e.g., approximately 65 million unique proteins for ESM-2 [27]), PLMs can encode proteins into representations that encompass rich protein-related information, including structures, functional properties, and evolutionary relationships. At a higher level, DNA foundation models can be adapted to larger molecules, such as bacteria and plasmids, by treating each nucleotide as a token. For example, the genomic foundation model, Evo [28], effectively extends single-nucleotide DNA modeling to a whole-genome scale, featuring a context length of 650 kilobases. However, to date, there are no multi-modal algorithms for DNA-text learning developed based on Evo. Furthermore, although the scientific LLM Galactica [29] included 0.1 billion nucleotides from human genomes in its training corpus by combining nucleotide tokens with textual tokens into the same vocabulary, it lacks knowledge related to microbes.

## Material and Methods

### Overview and Design Rationale of PlasRAG

Fig. 2 sketches the two main modules of PlasRAG: (1) sequence-to-text (S2T) plasmid characterization, and (2) text-to-sequence (T2S) plasmid retrieval. The two modules share a core multi-modal IR model [30] designed to learn the associations between plasmid DNA sequence features and property annotations expressed in natural language (e.g., *The plasmid encodes the WHO-prioritized ARG IMP-1*.). The input data for the IR model comprises sequence-text pairs, which are positively labeled if the plasmid can be annotated with the corresponding textual property, and negatively labeled otherwise. Within the IR model, each pair will first undergo two encoders: one to obtain a DNA sequence embedding that encodes plasmid functions and properties, and another to produce a text embedding that encodes biomedical knowledge. Subsequently, the two embeddings will be linearly projected into a shared cross-modal embedding space, where their cosine similarity will be calculated. A higher similarity indicates a greater probability that the cross-modal pair is positive. Thus, during training, the IR model will optimize the similarities to align with the gold labels of our curated sequence-text pair training set.

**Fig. 2.**
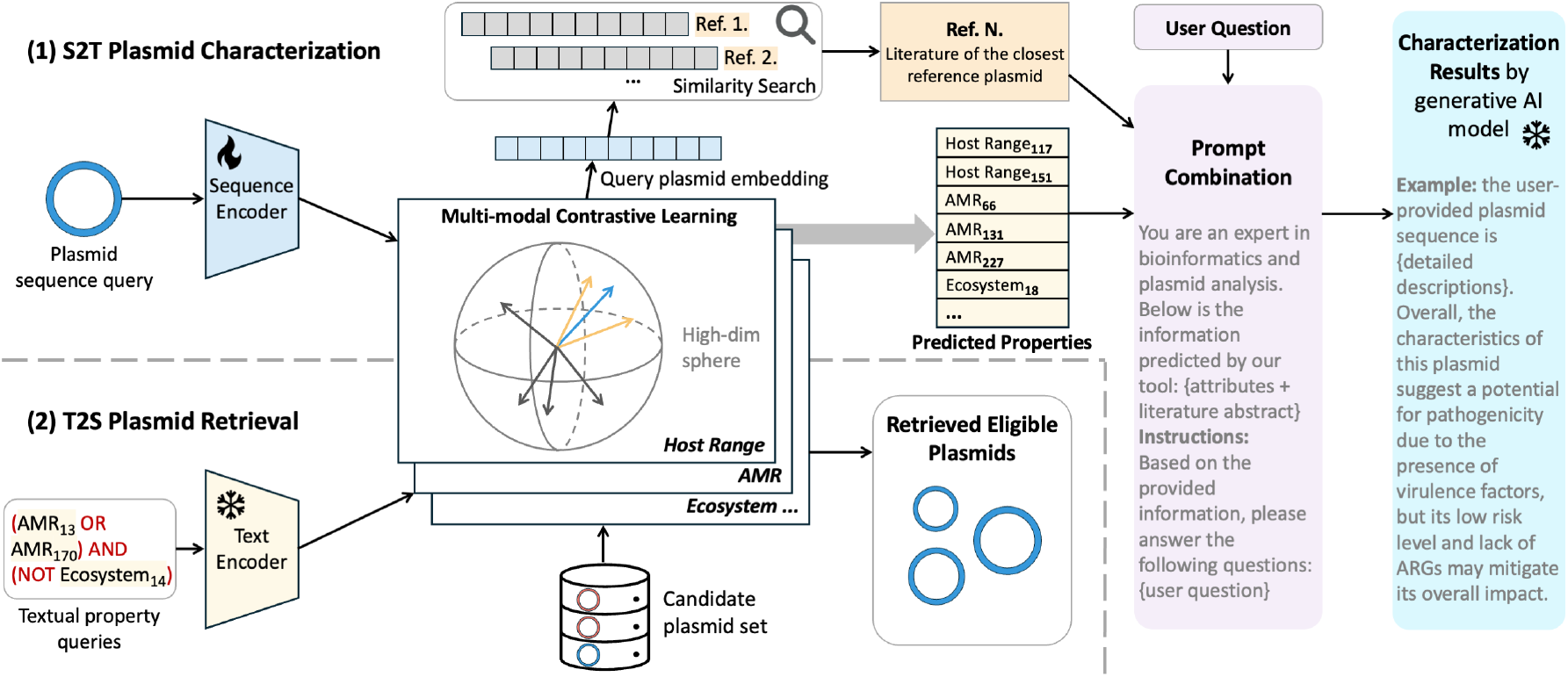
The workflow of PlasRAG’s two modules: (1) sequence-to-text (S2T) plasmid characterization and (2) text-to-sequence (T2S) plasmid retrieval, which are separated by a gray dashed line. Due to space limitations, we only present 3 out of the 10 multi-modal IR models shared by the two modules in the center of the figure. Additionally, each textual property (a complete sentence) is abbreviated in the format of Facet_index_ and displayed with a yellow background. For example, AMR_13_ represents the 13th text in the *AMR* sub-vocabulary. The fire symbol indicates that the sequence encoder is involved in training, while the two snowflake symbols signify that the parameters of the text encoder and the generative model remain frozen during training. (1) The input is a plasmid DNA sequence (represented by a blue circle), and the characterization result is the output generated by the LLM using the RAG framework. (2) Users need to select the interested property texts to create a query condition (Boolean expression) and optionally specify a candidate plasmid set for retrieval. The retrieval results include eligible plasmids that satisfy the query condition.

To enhance PlasRAG’s characterization ability, we define 10 property facets highly relevant to plasmid research, including *AMR, Host Range, Ecosystem, Ecological Host, Fundamental Property, HMR, Incompatibility (Inc) Group, Mobility, Risk Index*, and *VF*. A fixed vocabulary consisting of complete descriptive sentences is defined for each facet, with more detailed vocabularies available in Supplementary Table S2. In addition, one single IR model will be trained for each facet. Thus, PlasRAG contains 10 sub-models with identical architecture but different parameters.

### Sequence-to-text (S2T) plasmid characterization

To utilize this module, users can input their plasmid sequences of interest as queries, which will be forwarded to the IR models to align with all pre-indexed embeddings of the 10-faceted textual properties. As shown in Fig. 2(1), all texts with a similarity to the query plasmid greater than the threshold *θ* will be retained as predicted properties. On the other hand, an overall embedding for the query plasmid will also be computed by synthesizing the sequence embeddings from the 10 sub-models, as detailed in the following section, ‘Sequence-Text Alignment’. This overall query embedding reflects the functionality and properties of the query plasmid from a comprehensive perspective and plays a crucial role in the literature recommendation phase. Specifically, by comparing it to our pre-indexed reference plasmid embedding database, where each reference plasmid is linked to relevant literature, we can identify the closest reference plasmid with the smallest Euclidean distance to the query embedding. Finally, given the three components, namely predicted properties, the most relevant literature, and the user’s question about the query plasmid, a flexible template will combine them into a coherent prompt. The powerful, instruction-tuned LLM, Llama-3 [31], is utilized for interacting with users on plasmid analysis, with a low temperature parameter of *T* = 0.2 specified for more reliable generation.

### Text-to-sequence (T2S) plasmid retrieval

As depicted in Fig. 2(2), the retrieval module is designed to filter eligible plasmids from either our default plasmid database or a user-provided set of candidate plasmids. To implement this module, users need to select desired property texts from the 10-faceted vocabulary and combine them using logical operators (AND, OR, NOT) to form a query condition. When processing a candidate plasmid, each text in the query condition can be treated as a Boolean expression that evaluates to True if the corresponding sequence-text similarity (predicted by the IR models) exceeds the threshold *θ*, and to False otherwise. Therefore, the final retrieval result comprises all eligible plasmids that make the overall query condition evaluate to True.

In the following sections, we will illustrate each key aspect of the PlasRAG method, including the text and sequence encoders, the multi-modal contrastive learning algorithm, and the details of data curation and model training, step by step.

### Encoding Property Texts: the MedCPT Encoder

To enable the text encoder to effectively extract specialized knowledge from plasmid properties, we expand short phrases annotated from the raw databases [9, 10, 12] into complete sentences. For example, the VF property *type 3 fimbriae* is expanded to: *The plasmid carries the type 3 fimbriae virulence factor, which belongs to the biofilm category*. Another example is the host taxonomy *family Enterobacteriaceae*, which is elaborated as: *The plasmid is hosted by bacteria belonging to the Enterobacteriaceae family*. To encode these sentences into meaningful text embeddings, we employ MedCPT [26], the SOTA LM designed for biomedical IR, as introduced in the ‘Related work’ section. MedCPT consists of three BERT[22]-based sub-models, all initialized from the BioMedBERT_110M_ base model [25]: one for encoding short-text user queries, one for encoding long-text literature abstracts, and one for ranking the retrieval results. Among these, we select the short-text query encoder (QEnc) because it specializes in generating semantically rich embeddings of short biomedical sentences, aligning well with our expanded property texts. Specifically, for each expanded sentence *t*, it is encoded into the text embedding *T*_*t*_ by the QEnc:

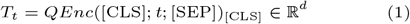

where [CLS] and [SEP] are special tokens defined in the original BERT. The text embedding *T*_*t*_ is derived from the last hidden state of QEnc corresponding to the [CLS] token. Here, *d* = 768 denotes the hidden size of the BERT-based QEnc model. To evaluate the effectiveness of the resulting text embedding *T*_*t*_, we select three representative VF categories from the bacterial virulence factor reference database, VFDB [32]. Subsequently, the corresponding *T*_*t*_ are generated by QEnc for all VF texts across the three selected categories. As shown in Fig. 3, we visualize all the examined VF embeddings *T*_*t*_ using the principal component analysis (PCA) method [33]. The clear distinction of embeddings among the three VF categories demonstrates that QEnc effectively captures the semantic similarities between biomedical sentences, particularly those related to plasmid properties. Notably, during the training phase, all text embeddings are pre-computed and stored, while the QEnc parameters remain fixed.

**Fig. 3.**
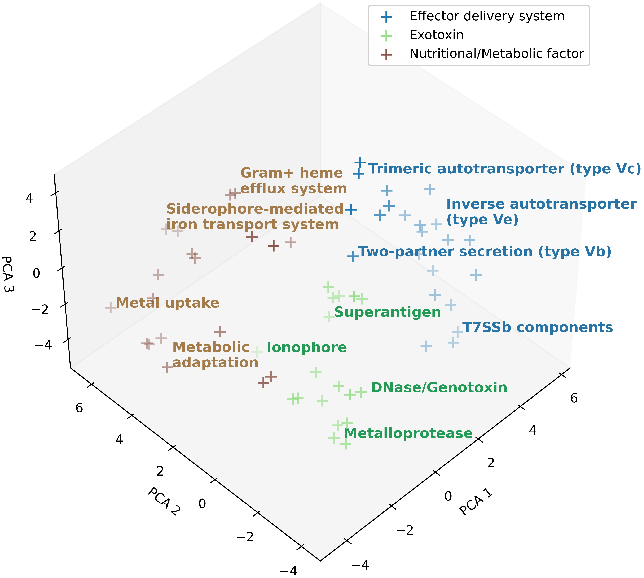
Three-dimensional (3D) visualization of encoded short-text embeddings by the QEnc model for all VFs across three selected categories: *Effector Delivery System, Exotoxin*, and *Nutritional/Metabolic Factor*. Each plus sign represents the visualized text embedding for a VF, with some representatives labeled by their descriptive texts surrounding the plus signs. We observed that the MedCPT QEnc model effectively captures the functional semantics of VF sentences.

### Encoding DNA Sequences: Cross-attention-based Perceiver Resampler

We construct the 10-faceted vocabulary for PlasRAG to encompass plasmid properties as comprehensively as possible. As a result, even within a sub-model of a single facet, there are property texts that characterize plasmids from various perspectives. For example, within the *Fundamental Property* model, we curate plasmid property texts that encompass completeness, topology, data source, gram-staining reaction of bacterial hosts, and phage-plasmid identification. Therefore, to improve performance in capturing the complex sequence-text associations, we design the PlasRAG sequence encoder to produce multiple sub-embeddings for each plasmid, in contrast to the traditional encoder that learns a single embedding. For a sub-model that includes *N* property texts (e.g., *N* = 205 for the *VF* model), the PlasRAG sequence encoder generates *N* sub-embeddings for each plasmid, enabling a more informative one-to-one alignment with the vocabulary. In the subsequent ‘Ablation Studies’ section, we will validate this unique design by comparing it with other sequence encoders based on traditional DL models. Specifically, the encoder architecture is implemented by connecting a cross-attention-based Perceiver Resampler model [34] to the SOTA protein language model (PLM) ESM-2 [27].

Given the strong associations between plasmid functionality, their properties, and the encoded proteins [35], we choose to target the coding regions (CDSs) of plasmids for feature extraction. As shown in Fig. 4, Prodigal [36] is first used for gene recognition and translation of the plasmid DNA sequence. The resulting proteins are then input individually into ESM-2 (with frozen parameters) to generate biologically meaningful embeddings that capture rich features, particularly the relationship between the 1D protein sequences and their corresponding 3D structures. The embedding of each protein is calculated from the global average pooling of the per-residue embeddings of all its amino acids (AAs). Following the encoding order of the proteins in the plasmid, we combine their ESM-2 embeddings as spatial features and pass them to the Perceiver Resampler, which transforms the varying-sized protein features into fixed-sized sub-embeddings for subsequent DNA sequence-text alignment.

**Fig. 4.**
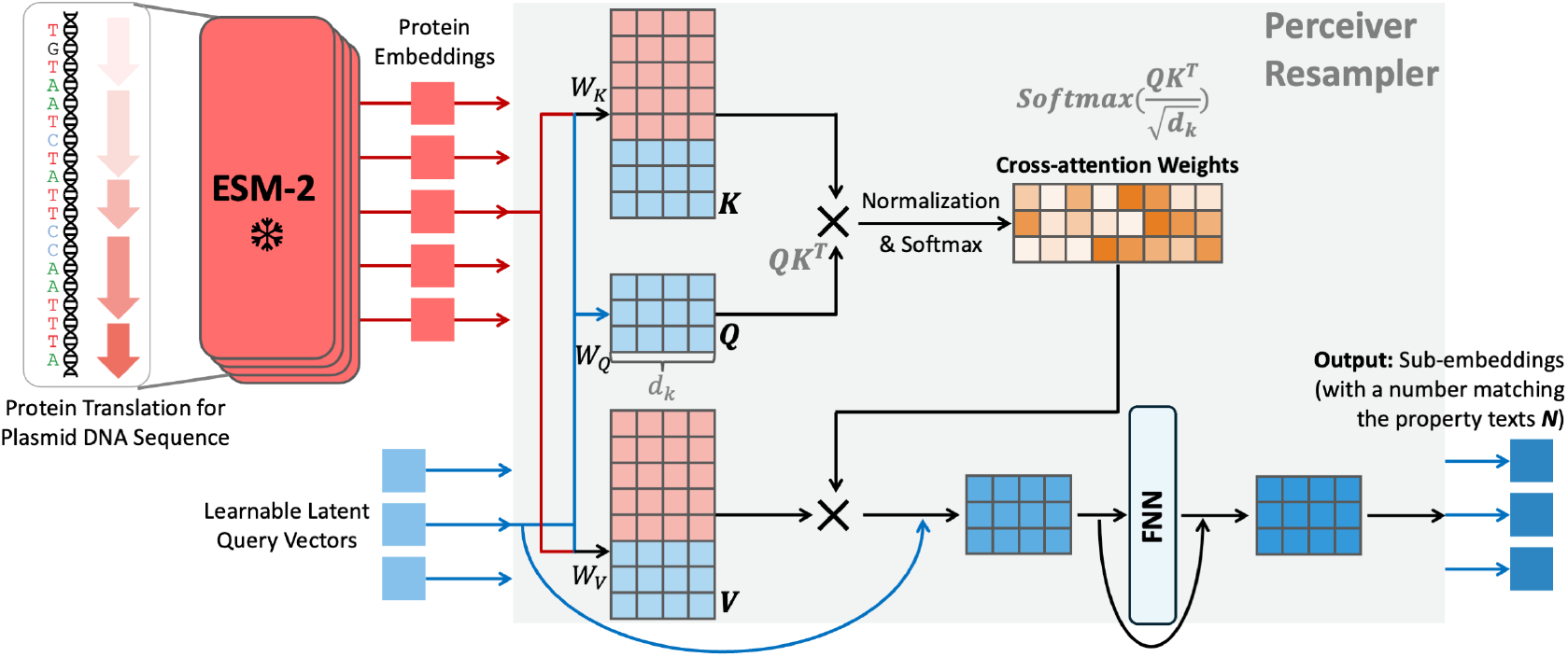
The model architecture of PlasRAG’s sequence encoder, which integrates the PLM ESM-2 with the Perceiver Resampler. The five red arrows in gradient colors represent translated proteins from the plasmid sequence. Each square, whether input or output of the Perceiver Resampler, signifies an embedding of dimension *d* = 768. To illustrate the Perceiver Resampler model, we use a toy example with *len* = 5, *N* = 3, and *d*_*k*_ = 4. A simple example is the three texts in the *Mobility* facet: ‘Mobilizable plasmid.’, ‘Non-mobilizable plasmid.’, and ‘Conjugative plasmid.’ Additionally, the two curved arrows indicate the two residual connections.

As depicted in Fig. 4, the core mechanism of the Perceiver Resampler is to calculate cross-attention [20] between the combined protein embeddings *E*_*prot*_ *∈* ℝ^*len×d*^ and a learnable latent query matrix *X ∈* ℝ^*N×d*^. Here, *len* = 300 denotes the maximum number of proteins encoded in a plasmid. For longer plasmids, only the first 300 proteins are considered, while shorter plasmids are padded at the end using a padding token. *d* = 768 represents the dimension of the protein embeddings, which is equal to the hidden size of the Perceiver Resampler’s Transformer encoder. Additionally, *N* represents the number of property texts curated for a sub-model of a facet. The calculation of cross-attention can be expressed as follows:

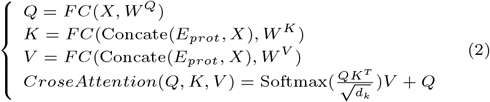

where the queries 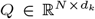 are derived from the learnable latent queries *X*, while the keys 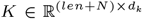 and values 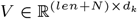 are obtained from the concatenation of *E*_*prot*_ and *X*. The projection matrices 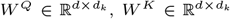, and 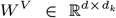 represent the learned parameters in the fully connected (FC) layers for *Q, K*, and *V*, respectively. The first residual connection (RC) [37] is then applied to the *CrossAttention* by adding the queries *Q*. Subsequently, *CrossAttention* passes through a feed-forward network, which incorporates the second RC, to produce the final output *S* of the entire sequence encoder. The output *S ∈* ℝ^*N×d*^ represents the multiple sub-embeddings extracted for a single plasmid. In the next section, we will elaborate on multi-modal contrastive learning to achieve sequence-text alignment between the text and sequence embeddings.

### Sequence-Text Alignment: Multi-modal Contrastive Learning

After encoding the text and sequence embeddings, a multimodal contrastive learning algorithm is employed to learn the intricate sequence-text correlations. The two-modality embeddings are linearly projected into a shared multi-modal embedding space, where their similarities are optimized under supervision from the gold labels of our curated training set. To illustrate more specific calculations, we present a simplified example in Fig. 5, featuring one representative plasmid *p* and a toy property vocabulary from the *Host Range* model. Given the embeddings of *N* in-vocabulary texts *T ∈* ℝ^*N×d*^ and the multiple sub-embeddings learned for plasmid *p S ∈* ℝ^*N×d*^ of the same dimension, we calculate the row-wise cosine similarity between them as follows:

**Fig. 5.**
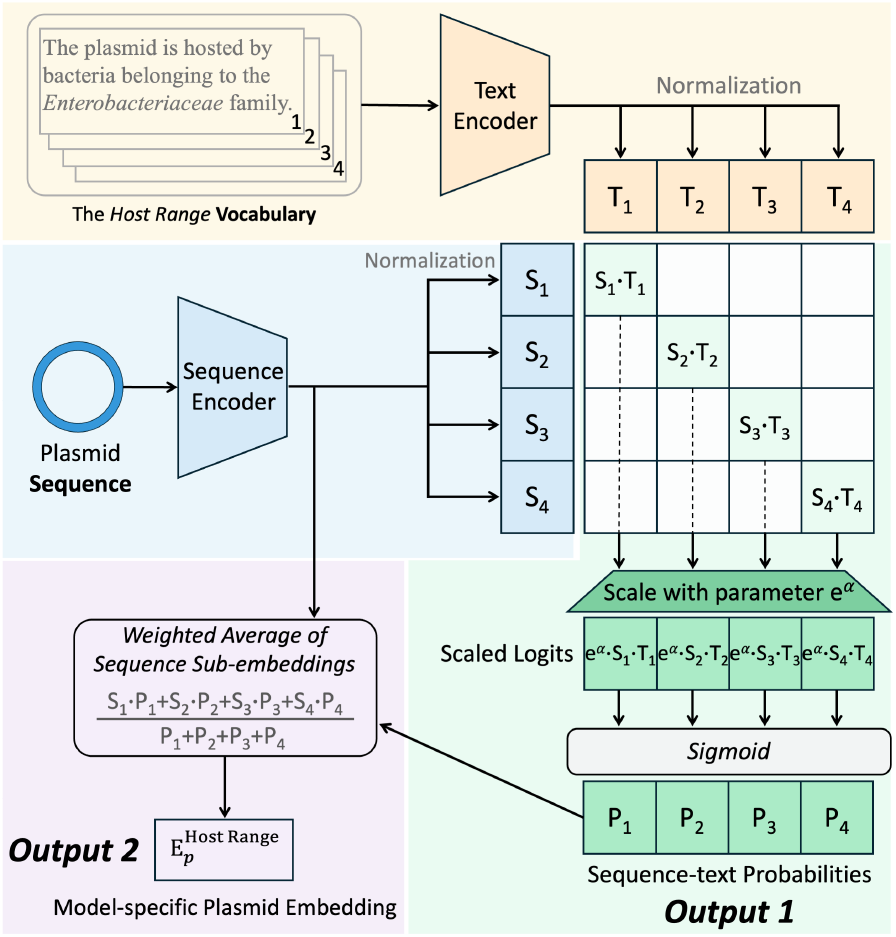
Illustration of the multi-modal contrastive learning algorithm using a toy example with *N* = 4 texts within the *Host Range* model. The figure is divided into four sections, each with a different colored background. Yellow for the text encoding phase, Blue for the sequence encoding phase, Green for the calculation of the sequence-text cosine probabilities, and Purple for the synthesis of the model-specific plasmid embedding 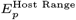.

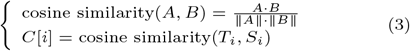

where *A ∈* ℝ and *B ∈* ℝ are two example vectors used to illustrate the calculation of cosine similarity. *T*_*i*_ and *S*_*i*_ (*I ∈* [0, *N*)) denote the *i*th row of *T* and *S* (aligning with the *i*th in-vocabulary text), respectively. As shown in Fig. 5, the resulting similarity vector *C ∈* ℝ^*N*^ corresponds to the diagonal of the square matrix *S · T*^*T*^ and will be retained for evaluating our defined sequence-text pair (STP) loss. However, the cosine similarity values in C range from −1 to 1, which restricts the sequence-text probabilities to approximately 0.269 to 0.731 when applying the sigmoid function (*σ*). Therefore, we scale the raw similarity vector *C* (logits) using a learnable temperature parameter *e*^*α*^ to expand the probability range. In summary, the STP loss (ℒ_*STP*_), a variant of the binary cross-entropy (BCE) loss, is formulated as follows:

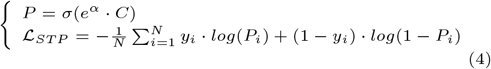

where *P ∈* ℝ^*N*^ represents the predicted sequence-text probabilities, and *P*_*i*_ denotes the probability that plasmid *p* can be annotated with the *i*th text. *y*_*i*_ is the *i*th gold label, taking the value 1 if plasmid *p* is annotated with the *i*th text in the training set, and 0 otherwise. As our training set is integrated from crowd-labeling across multiple plasmid databases, a special case may arise if the database curating plasmid *p* never annotates plasmids with the *i*th text. In this situation, *y*_*i*_ is masked with −100, and the loss calculation is skipped for the corresponding *p*-*i* sequence-text pair. By optimizing the multi-modal IR models, particularly the sequence encoder, towards the STP loss, the goal is to maximize the similarity between correct plasmid sequence-text pairs while minimizing the similarity for incorrect pairs.

Another key objective of the multi-modal IR models is to synthesize the overall embedding for plasmid *p*, which facilitates the following literature recommendation. At a higher level, a model-specific plasmid embedding 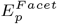 will be synthesized individually for each of the 10 sub-models, and a weight vector of hyperparameters *W ∈* ℝ^10^ will average the 10 model-specific embeddings into the final overall embedding *E*_*p*_. Fig. 5 illustrates the calculation of the overall plasmid embedding 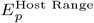, specifically by averaging the sub-embeddings *S* using *P* as weights. In this way, the overall plasmid embedding *E*_*p*_ will encode rich features of plasmid *p*’s functionality and properties from various important perspectives.

### Data Curation and Model Training

#### Plasmid Sequence Data

We collected plasmid sequences from three high-quality plasmid databases: PIPdb [10], IMG/PR [9], and PLSDB [12], to train the PlasRAG models. We exclude megaplasmids, which are typically large plasmids exceeding 250 kbp in length, due to their close relationship with chromids (secondary chromosomes) [38]. As a result, a total of 1,521,635 qualified plasmids remain. To rigorously evaluate PlasRAG’s performance on novel plasmids, we allocate 100,000 plasmids to the validation and test sets following the plasmid segment cluster (PSC) rules defined by PIPdb. Specifically, all plasmids within the same PSC that share an average nucleotide identity (ANI) above 76% are collectively assigned to one of the training, validation, or test sets, ensuring sufficient dissimilarity among the three partitions. Further details regarding the dataset construction are provided in Table 1. Subsequently, we use Prodigal [36] to translate the genes of all plasmids, yielding a total of 37,050,904 plasmid-encoded proteins. Although the protein embeddings will be pre-computed and won’t increase training costs, they will still need to be preloaded into CPU memory, resulting in significant resource consumption. For instance, even though we chose the lightweight ESM-2_650M_ model to encode proteins into vectors of dimension 1,280, approximately 176.7 GB of CPU memory is required. Thus, we cluster the raw translated proteins according to the UniRef90 cutoff standard [39] using the MMseqs2 algorithm [40] and select a representative for each cluster. This process produces 4,925,205 deduplicated proteins, reducing CPU memory consumption to 23.5 GB.

**Table 1.**
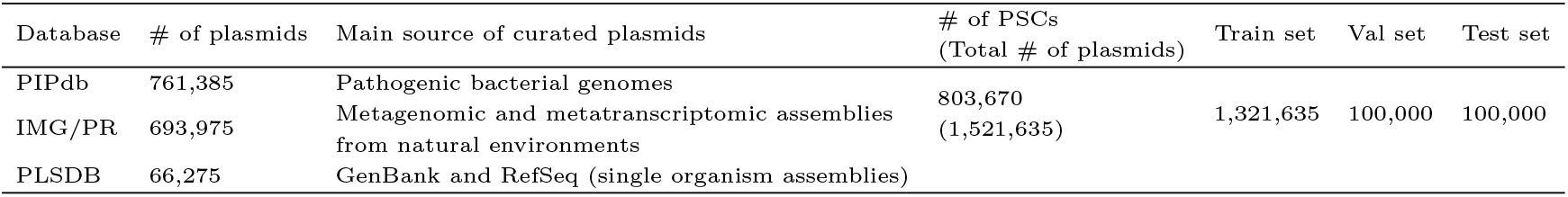
Detailed information about the three well-curated plasmid databases and the dataset we constructed.

#### Property Text Data

We manually collect raw property annotations from the three databases to curate the plasmid sequence-text pair dataset for our defined 10 facets. Following the method described in the ‘Encoding Property Texts’ section, all collected raw properties are expanded into complete descriptive sentences, preparing for the subsequent text embedding with the MedCPT model. The detailed information listing which database annotates plasmids from which facets, along with brief descriptions of their annotation contents, is shown in Supplementary Table S1. This table also displays the value of the text number *N* for each facet. We can take a closer look at the *Ecological Host* facet to illustrate the missing annotation condition more clearly. For plasmids sourced from PIPdb or IMG/PR, there is no annotated ecological host information. In this case, the masked label vector filled with −100 will be assigned to these plasmids. The complete version of our constructed 10-faceted vocabulary is available in Supplementary Table S2.

When processing the plasmid sequence-text pairs, two property facets, *Host Range* and *Risk Index*, require additional procedures. First, due to the prevalence of broad-host-range (BHR) plasmids [41], we use the concept of plasmid taxonomic units (PTUs) [42] to determine the plasmid host range. In detail, for all plasmids within the same PTU, their host range labels are assigned as the lowest common ancestor (LCA) of their host species in the phylogenetic tree. Second, we determine four plasmid risk levels [10], ranging from *Minimal* to *Critical*, for each risk category evaluated based on various factors, such as the number of insertion sequences harbored by the plasmid. Instead of treating the four levels as distinct property texts, we consider them as a single text and assign each risk level a floating soft label during training. For instance, a soft label of 0.4 is assigned to the *Moderate* risk, while 1.0 is assigned to the *Critical* risk. The final step is to curate the reference plasmids for literature recommendations. Specifically, all plasmids with annotated PubMed IDs in the RefSeq [43] database are retained as reference plasmids, and their associated literature abstracts and citation formats are retrieved using the Entrez Fetch tool [44]. Consequently, 10,282 reference plasmids have been cataloged and linked to 3,120 biomedical publications.

#### Model Training

Before training PlasRAG, the protein and text embeddings are pre-computed and stored locally using the ESM-2_650M_ and the MedCPT QEnc_110M_ model. Next, we train 10 sub-models, each corresponding to one of the 10 property facets. As mentioned in previous sections, only the Perceiver Resampler (within the sequence encoder) is involved in parameter optimization, which makes the PlasRAG models highly efficient for training. In total, the 10 sub-models comprise 89.3 million parameters, allowing us to accelerate the training process with a larger batch size. Specifically, we conduct the model training with a learning rate of 2e-4 and a batch size of 2,048 for 20 epochs on a single NVIDIA A100 80GB graphics card. The best models are saved based on evaluations using the validation set after each epoch. A warm-up rate of 0.1 is applied, causing the learning rate to increase linearly for the first 2 epochs and then decay linearly thereafter. In addition, mixed precision training [45] is employed for faster computations by storing activations (output values produced by neurons) and computing gradients in half-precision (FP16) while storing other components (gradients, model weights, and losses) in full-precision (FP32). In summary, the PlasRAG models are resource-efficient during both the training and prediction phases, with a detailed evaluation of computational consumption provided in Supplementary Table S3.

## Results

### Experimental Setup

#### Metrics

We employ two metrics commonly used in information retrieval (IR) systems: F1-score and Precision at *k* (*P* @*k*), to evaluate the performance of PlasRAG’s core sequence-text IR models. Since a single IR model can symmetrically retrieve plasmids based on query texts (text-centric) and retrieve texts based on query plasmids (sequence-centric), strong performance in the text-centric approach also indicates effective performance in the sequence-centric approach. Therefore, we choose the text-centric evaluation for the 10-faceted IR models in this work. First, the F1-score is calculated individually for each text *t* and then weighted by the number of testing plasmids *w*_*t*_ truly annotated by *t* (gold labels), as detailed in Equation 5.

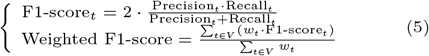

where *V* is the vocabulary consisting of *N* texts for a single property facet, and the performance for each facet will be evaluated independently.

*P* @*k* is a specialized metric for evaluating the ranking and recommendation capabilities of an IR system. Specifically, *P* @*k* is determined as the proportion of testing plasmids truly annotated by text *t* among the top *k* predicted plasmids with the highest scores (based on cosine similarity with text *t*), as defined in Equation 6.

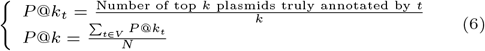

Considering the scale of the curated test set (100,000 testing plasmids), we choose *k* = 1000 to evaluate our IR model.

#### The PlasX Plasmid System (PS) Dataset

Yu et al. designed a machine-learning method, PlasX [46], to identify 68,350 novel plasmids from 1,782 human gut metagenomes across 16 countries, which include 5 non-industrialized and 11 industrialized nations. These newly discovered plasmids were further organized into 1,169 defined evolutionarily cohesive plasmid systems (PSs). The resulting PS dataset serves as a valuable resource for evaluating PlasRAG’s ability to characterize novel plasmids with high-quality origin and evolution information. Members of the same PS can be classified into backbone and compound plasmids. Backbone plasmids are the fundamental components or subsequences of compound plasmids, encoding only the genes essential for the housekeeping functions of plasmids, such as maintenance and transfer. In contrast, compound plasmids encode additional cargo genes, such as ARGs, based on backbone plasmids. This division aligns with the observation that the same backbone frequently recurs during plasmid evolution [47], while different cargo genes emerge in response to various environmental pressures. Additionally, the country information associated with each PS allows us to identify ecological patterns of plasmids derived from humans with varying lifestyles.

#### Ablation Studies: Evaluating Design Rationale

In this section, we conducted ablation studies on PlasRAG’s multi-modal model architecture, focusing on two key aspects: the sequence and text encoders. To effectively evaluate the two encoders, we adhered to the principle of changing only one component at a time. Fig. 6 illustrates the nine experimental combinations: one from the original PlasRAG, four that replace the sequence encoder, and four that substitute the text encoder. All model combinations were trained using the same dataset detailed in Table 1 and the same parameters, including the learning rate.

**Fig. 6.**
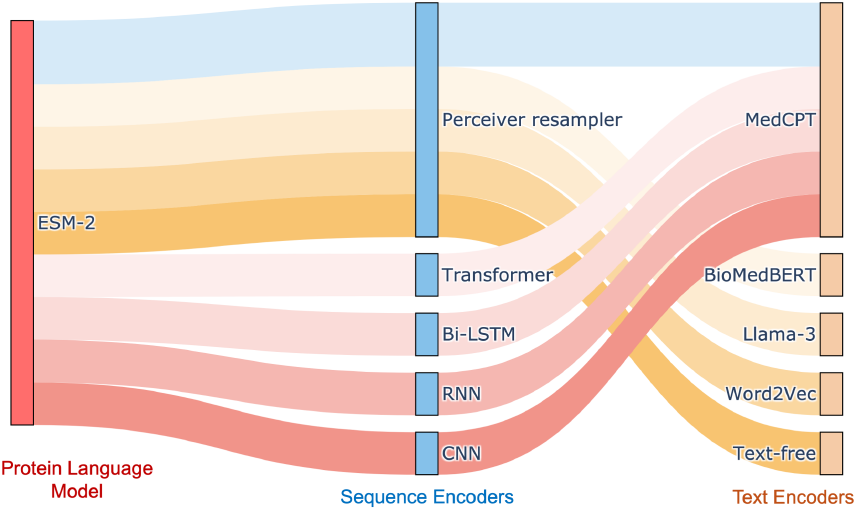
The Sankey diagram illustrating the nine encoder combinations created for the ablation studies. We aim to evaluate the effectiveness of PlasRAG’s two encoders: the Perceiver Resampler (sequence encoder) and the MedCPT QEnc (text encoder). Notably, we retained the same ESM-2 model for raw sequence feature extraction, as it has been extensively validated to demonstrate strong performance in protein understanding. In the figure, each blue bar represents a candidate sequence encoder connected to ESM-2, while each yellow bar represents a candidate text encoder for encoding property texts. The top blue link indicates the default setup for PlasRAG, the yellow links represent combinations for evaluating candidate text encoders, and the red links denote combinations for assessing candidate sequence encoders.

#### Design of the ablation studies

To test sequence encoders, we compared the default Perceiver Resampler with four classical DL models: Transformer [20], Bidirectional Long Short-Term Memory (Bi-LSTM), Recurrent

Neural Network (RNN), and Convolutional Neural Network (CNN). The foundational ESM-2 model remained unchanged for all experimental combinations. Thus, the input to the five benchmarked sequence encoders consists of sequential ESM-2 embeddings in the same order as the proteins encoded within the plasmid. The first three models are specialized in capturing features from sequential data, making them particularly well-suited for encoding plasmids. Conversely, the CNN model is originally designed for grid-like data, such as images. Therefore, we adapted the CNN to encode plasmids by inputting vertically concatenated ESM-2 protein embeddings in matrix form. For the text encoder testing, we replaced the default MedCPT QEnc model with two LMs: BioMedBERT [25] and Llama-3 [31], in addition to the Word2Vec word embedding method. We also incorporated a text-free approach that does not utilize any textual features, defining text retrieval as a traditional multi-label classification problem based solely on sequence features. Finally, we compared our implemented DL models with the alignment-based tool BLASTN [48] by aligning test plasmids with training plasmids and determining the predicted properties based on the best-hit plasmids. The benchmark results are presented in Table 2 for the F1-score and in Table 3 for the P@1000 metric. We observed that PlasRAG’s original design achieved the overall best performance. The lower performance of BLASTN aligns with the challenge we previously presented, where alignment-based methods struggle particularly to characterize novel plasmids that exhibit low similarity to known plasmids.

**Table 2.** Results of the ablation experiment across 10 property facets on the plasmid test set, evaluated using the weighted F1-score metric.

| Candidate Components |  | Weighted Text-Centric F1-score |  |  |  |  |  |  |  |  |  |
| --- | --- | --- | --- | --- | --- | --- | --- | --- | --- | --- | --- |
| Sequence Encoder | Text Encoder | *AMR* | *Host Range* | *Ecological Host* | *Ecosystem* | *Basic Property* | *HMR* | *Inc Group* | *Mobility* | *Risk Index* | *VF* |
| **Perceiver Resampler** | **MedCPT (original)** | **94.0** | **83.1** | **81.5** | **77.7** | **92.9** | **96.5** | **96.2** | **97.4** | **83.7** | **90.7** |
| Transformer |  | 81.1 | 78.5 | 75.9 | 74.6 | 91.8 | 60.0 | 84.8 | 91.6 | 74.0 | 44.5 |
| Bi-LSTM |  | 91.0 | 81.0 | 77.6 | 76.5 | 92.6 | 78.4 | 93.0 | 96.3 | 80.1 | 75.4 |
| RNN |  | 91.1 | 80.1 | 77.0 | 76.6 | 92.4 | 77.8 | 92.7 | 96.0 | 78.8 | 70.7 |
| CNN |  | 90.9 | 79.9 | 76.5 | 76.2 | 92.0 | 78.6 | 92.1 | 96.1 | 78.4 | 72.1 |
| **Perceiver Resampler (original)** | BioMedBERT | 93.6 | 80.5 | 75.6 | 75.0 | 92.2 | 93.3 | 94.4 | 96.9 | 81.5 | 86.3 |
|  | Llama-3 | 93.3 | 79.8 | 74.7 | 74.3 | 92.0 | 93.0 | 93.9 | 96.8 | 80.7 | 85.7 |
|  | Word2Vec | 92.7 | 78.8 | 74.0 | 73.4 | 91.9 | 91.1 | 93.9 | 96.6 | 80.2 | 79.6 |
|  | Text-free | 91.7 | 76.7 | 72.8 | 74.7 | 92.4 | 68.4 | 90.0 | 96.6 | 80.5 | 19.7 |
| Alignment - BLASTN |  | 55.6 | 83.0 | 70.4 | 76.4 | 87.6 | 53.4 | 64.1 | 77.6 | 76.5 | 57.9 |

**Table 3.** Results of the ablation experiment across 10 property facets on the plasmid test set, evaluated using the P@1000 metric.

| Candidate Components |  | Text-Centric P@1000 |  |  |  |  |  |  |  |  |  |
| --- | --- | --- | --- | --- | --- | --- | --- | --- | --- | --- | --- |
| Sequence Encoder | Text Encoder | *AMR* | *Host Range* | *Ecological Host* | *Ecosystem* | *Basic Property* | *HMR* | *Inc Group* | *Mobility* | *Risk Index* | *VF* |
| **Perceiver Resampler** | **MedCPT (original)** | **96.4** | **84.9** | **79.2** | **74.7** | **91.7** | **97.5** | **96.4** | **97.7** | **83.6** | **93.0** |
| Transformer |  | 84.2 | 74.3 | 73.0 | 68.6 | 88.2 | 62.9 | 87.0 | 88.6 | 56.9 | 49.9 |
| Bi-LSTM |  | 93.1 | 80.2 | 76.2 | 71.9 | 90.7 | 82.3 | 93.9 | 95.6 | 76.7 | 80.4 |
| RNN |  | 93.0 | 78.7 | 75.7 | 71.6 | 90.5 | 81.3 | 93.2 | 95.0 | 73.0 | 77.0 |
| CNN |  | 93.0 | 78.5 | 76.1 | 71.6 | 90.1 | 83.5 | 93.2 | 95.3 | 74.0 | 81.0 |
| **Perceiver Resampler (original)** | BioMedBERT | 95.8 | 80.6 | 74.7 | 70.8 | 90.9 | 94.7 | 95.1 | 96.9 | 80.0 | 88.6 |
|  | Llama-3 | 95.4 | 79.0 | 74.0 | 70.0 | 90.3 | 94.4 | 94.4 | 96.8 | 77.8 | 87.9 |
|  | Word2Vec | 94.9 | 77.1 | 72.9 | 68.3 | 90.1 | 92.6 | 94.2 | 96.5 | 78.1 | 82.6 |
|  | Text-free | 94.1 | 73.4 | 70.2 | 71.1 | 90.1 | 76.8 | 91.2 | 96.0 | 78.5 | 28.6 |
| Alignment - BLASTN |  | 49.5 | 79.9 | 75.2 | 71.2 | 75.7 | 37.3 | 62.8 | 67.3 | 62.8 | 45.2 |

#### Insights from the sequence encoder benchmark

Three insights arise from the sequence encoder comparisons. First, the Perceiver Resampler achieved the best performance across all 10-faceted retrieval tasks, indicating its enhanced ability to integrate multi-modal data by transforming raw features into fixed-size embeddings that align with text. Second, the Perceiver Resampler notably outperforms other candidates in property facets that include more texts, such as HMR and VF. This advantage stems from the learned multiple sub-embeddings for each plasmid, allowing for one unique sub-embedding to be aligned with each text embedding. Along with the improvement in sequence feature expression, the imbalances in plasmids annotated by each text can also be mitigated. Third, it is somewhat counterintuitive that the Transformer did not perform optimally, despite being an SOTA model adept at capturing long-distance dependencies. One possible reason is its reduced sensitivity to the order of the proteins encoded in plasmids, which is crucial for modeling plasmid sequences.

#### Insights from the text encoder benchmark

The superior overall performance of the MedCPT QEnc model indicates that the MedCPT embeddings effectively convey valuable insights from understanding plasmid property texts. Four additional insights are worth noting. First, the original BioMedBERT demonstrated a weaker capability to extract specialized information from plasmid-related texts compared to MedCPT. This suggests that continual adaptation, such as the contrastive learning approach of MedCPT starting from BioMedBERT, is essential for optimizing downstream tasks. Second, while Llama-3 is an excellent generative LLM for various NLP tasks, such as conversation, its performance on our tasks is hindered due to its unsuitability for text encoding and lack of emphasis on biomedical-specific knowledge. Third, the Word2Vec model excels at capturing syntactic embeddings for individual words. However, it achieves suboptimal performance in situations where a comprehensive understanding of a whole sentence is required. Finally, the poor performance of the text-free approach, especially regarding three text-rich facets: *Host Range, HMR*, and *VF*, demonstrated that our formulation of plasmid characterization as sequence-text multi-modal learning is valid.

#### Visualization of the Multi-modal Embedding Space

In this section, we utilized the PCA method to visualize the sequence-text embedding space, aiding in the interpretation of our multi-modal contrastive learning algorithm. As shown in Fig. 7, we presented three cases by measuring the sub-embeddings *S*, the overall plasmid embedding *E*_*p*_, and the text embeddings *T* (for more details, refer to the ‘Methods’ section). In Fig. 7A, we selected a representative text from the *Risk Index* facet and randomly chose 100 positive plasmids that were truly annotated by the text, along with 400 negative plasmids not associated with it, all from the test set. The results demonstrate a clear pattern: positive cosine similarity is observed for most embeddings of correct sequence-text pairs, while incorrect pairs consistently show negative similarity. This aligns closely with the STP loss objective based on cosine similarity optimization. Second, we initialized embeddings for six randomly selected texts, each from a different facet, along with a sample of their associated positive plasmids (Fig. 7B). We can observe that strong associations are learned for multi-faceted sequence-text pairs within a single embedding space, with the text embedding foundation provided by the same frozen MedCPT model. Finally, we evaluated the synthetic process of the overall plasmid embeddings *E*_p_. Specifically, 100 multidrug resistance (MDR) plasmids were randomly sampled for each of the top six most frequent antibiotic combinations (Fig. 7C). For instance, each orange circle in the figure represents the embedding 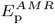 of a plasmid encoding ARGs that confer resistance to both Sulfonamide and Sulfone antibiotics. A clear distinction can be observed among plasmids with different combinations of antibiotic resistance. Thus, the synthesized embeddings effectively encode the comprehensive functionalities of plasmids, providing a robust foundation for searching for reference plasmids with associated literature.

**Fig. 7.**
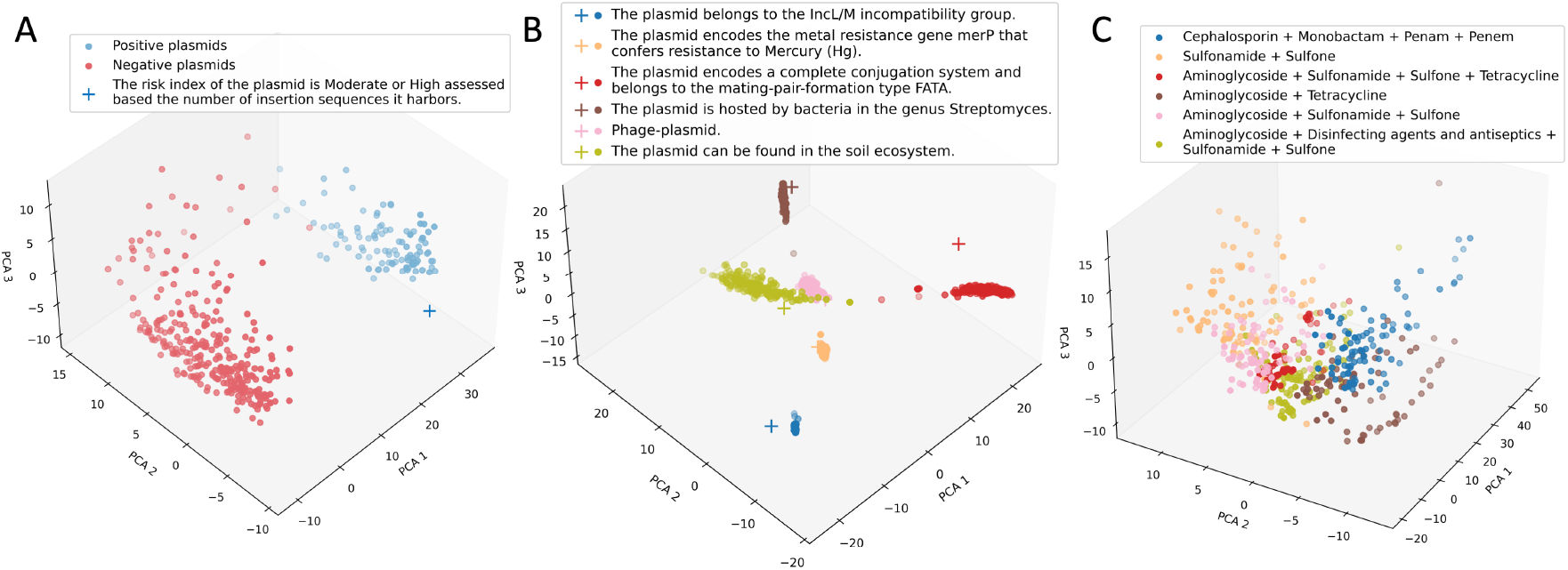
3D visualization of sequence and text embeddings illustrated through three cases. Each plus sign represents a text embedding produced by the MedCPT QEnc model, while each circle denotes an embedding learned from plasmid DNA sequences. (A) A text featuring both positive and negative plasmids to evaluate the multi-modal contrastive learning objective. A high similarity between the sequence and text embeddings indicates a greater probability that the plasmid is annotated by that text; (B) Correct sequence-text pairs from six different facets, demonstrating PlasRAG’s predictive capabilities across different plasmid property facets. Each color represents the positive plasmids and text associated with a single property; (C) Six groups of plasmids conferring resistance to different antibiotic combinations, assessing the effectiveness of the overall plasmid embedding synthesis. Each circle in (C) represents the overall plasmid embeddings averaged from multiple sub-embeddings.

### Specialized Comparison with a SOTA Tool in Mobility

PlasRAG is the first method designed to achieve our two defined plasmid-related goals: S2T characterization and T2S retrieval. Consequently, conducting exhaustive benchmarks across the 10 property facets is challenging due to the limited availability of plasmid characterization tools. To validate the accuracy of PlasRAG’s results, we conducted a targeted benchmark against MOBFinder [14], which is designed to characterize plasmids specifically within the *Mobility* facet. Specifically, both PlasRAG and MOBFinder can perform two tasks related to *Mobility*: plasmid mobility classification and relaxase type (mobilization type) prediction. The first task involves classifying a given plasmid into one of the three mobility types: non-mobilizable, mobilizable, or conjugative. The second task aims to predict the encoded MOB types (totaling 10 candidate types) for mobilizable and conjugative plasmids. The comparison results on the plasmid test set are shown in Table 4. We observed that PlasRAG nearly achieved perfect predictions for both tasks, while MOBFinder struggled to deliver accurate results. This demonstrates that the multi-modal IR model of PlasRAG remains reliable for unseen plasmid DNA, surpassing current tools in terms of practicality.

**Table 4.** Benchmark results comparing PlasRAG and MOBFinder evaluated on two *Mobility*-related tasks.

| Method | *Task* *Mobility Classification* |  | *Task* *Relaxase Type Prediction* |  |
| --- | --- | --- | --- | --- |
|  | F1-score | P@1000 | F1-score | P@1000 |
| PlasRAG | 97.7 | 99.9 | 96.8 | 97.8 |
| MOBFinder | 19.9 | 69.8 | 16.1 | 49.0 |

#### Case Studies: A Deep Dive into PlasRAG’s Functionality

*Template for the augmented prompt with the RAG pipeline* The upper section of Fig. 8 illustrates the RAG-based S2T plasmid characterization module designed within PlasRAG. Upon receiving a query plasmid, the retrieved knowledge, which includes predicted properties and recommended literature, will be integrated with user questions to augment the Llama-3 prompt. In Supplementary Section S3, we present the specific prompt template, organized into five key components: *Context, User Question, Predicted Properties, Literature*, and *Instruction*. The *Context* and *Instruction* sections are written as clear descriptions to provide the LLM with the task background and outline its objectives. Given that both components are essential, one is positioned at the beginning and the other at the end of the prompt, enhancing the LLM’s focus on them. In the *Instruction* section, we specify that the LLM should strictly utilize the provided information in the *Predicted Properties* and *Literature* sections, avoiding any additional details or hallucinations. With the augmented prompt, the LLM will produce more accurate and useful information for the query plasmid sequences, leveraging its capabilities in text understanding and reasoning.

**Fig. 8.**
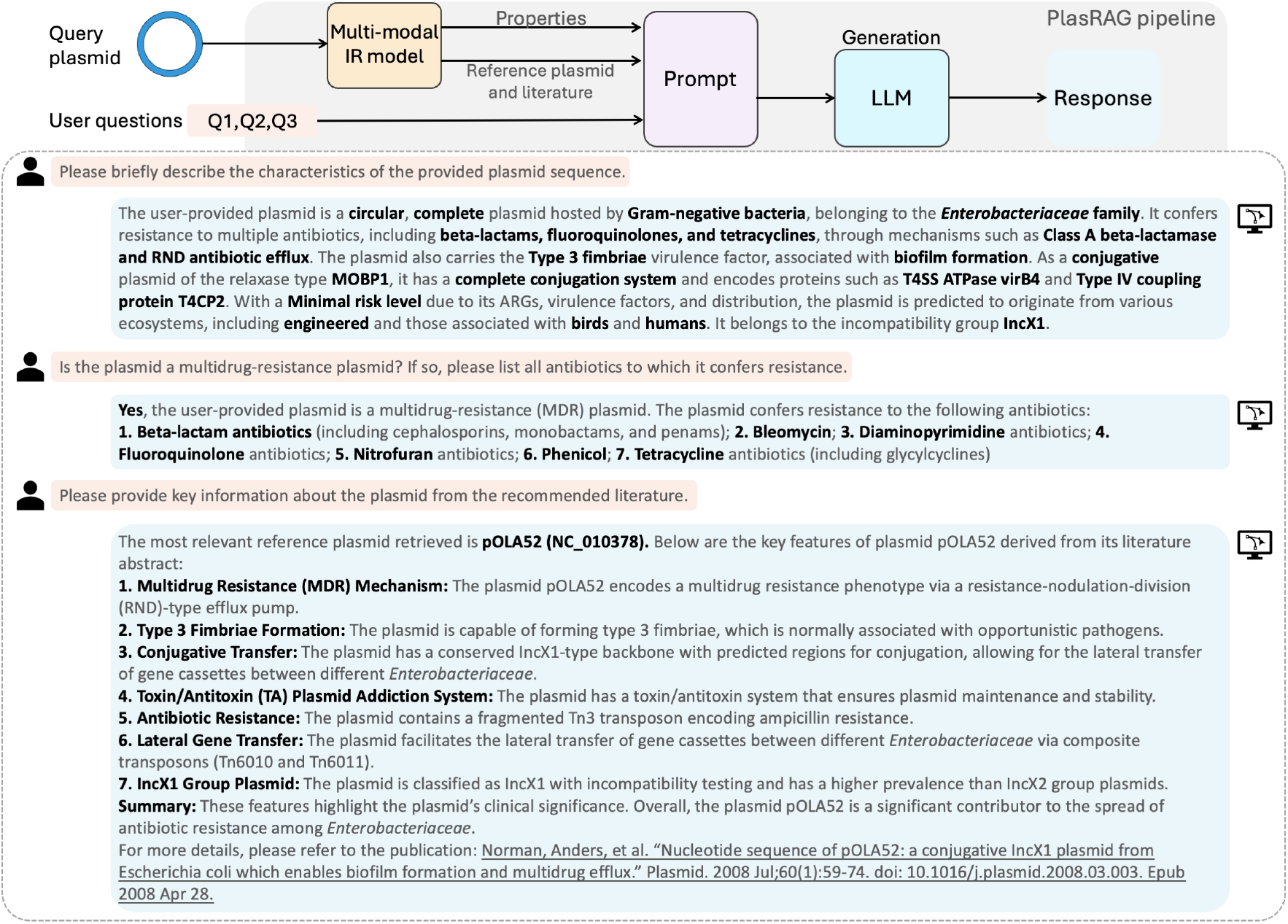
LLM generation instances for characterizing the plasmid pOLA52. The above pipeline illustrates the S2T plasmid characterization process, which incorporates the multi-modal IR model and the RAG framework. The conversational content in the box below represents instances of the interactive human–machine mode.

#### Case study of S2T plasmid characterization

In Fig. 8 and Supplementary Figure S1, we presented the PlasRAG characterization results for two plasmids: the reference plasmid pOLA52 (NC 010378), and a plasmid from the novel test set (*IMGPR plasmid 2772190775 000001*) identified from the *Clostridium argentinense* 89G isolate. Overall, the PlasRAG outputs can provide an accurate and comprehensive analysis of the query plasmids. More specifically, we noted a strong consistency in the descriptions summarized from our predicted properties (the first two responses) and the literature (the last response). This includes multi-resistance, type 3 fimbriae virulence, conjugation ability, host range within the family *Enterobacteriaceae*, and the IncX1 Inc group. In Supplementary Figure S2, we presented a detailed comparison between the LLM summary and the raw literature abstract. This demonstrated that, using our designed prompt template, Llama-3 can capture most key information while disregarding relatively less important details. Furthermore, the generated responses aligned closely with our retrieved knowledge and minimized hallucinations.

#### Case study of T2S plasmid retrieval

PlasRAG offers a comprehensive 10-faceted property vocabulary for formulating Boolean expression-based query conditions (Supplementary Table S2), and its retrieval performance has been demonstrated to be reliable in previous sections. To further illustrate the advantages of PlasRAG, we presented two cases for the T2S retrieval module in Supplementary Figures S3 and S4. The first case involved a provided candidate set of plasmids, while the second case retrieved eligible plasmids without any user submission, specifically selected from our pre-indexed database. Using the novel test set of 100,000 plasmids as the candidate set for evaluation, 487 eligible plasmids were retrieved based on a query condition combined from nine properties (Supplementary Figure S3). The evaluation yielded a recall of 96.2% and a precision of 91.6%, demonstrating the effectiveness of the T2S retrieval module on novel plasmids.

### Experiments on the Plasmid System (PS) Dataset

#### Initial evaluation of backbone and compound plasmids

In this section, we applied PlasRAG to the 1,169 plasmid systems (PSs) identified by Yu et al. in [46] to evaluate its capabilities in uncovering novel ecological insights. As detailed in the ‘Experimental Setup’ section, plasmids within a single PS can be categorized into backbone and compound plasmids. The former contains only conserved backbone sequences, while the latter encodes additional cargo genes, such as ARGs, based on the backbone plasmids within the same PS. To validate whether PlasRAG’s results align with this division scheme, we analyzed the differences in several facets of plasmid properties between the two groups. As shown in Fig. 9A, seven properties from five facets were selected as representatives, and the percentage of plasmids predicted with these properties in each PS is displayed as box plots to illustrate the distributions. In other words, the percentage value for each PS can be interpreted as its ‘property-specific purity’. The four properties with the highest overall purity values indicate that *Mobility, Host Range*, and *Basic Properties* are strongly dependent on plasmid backbone genes, particularly those responsible for replication, stability, and transfer. In contrast, the low purity values of *AMR* and *HMR* are expected, as these represent accessory functions encoded on plasmids that vary over time, influenced by the environmental pressures faced by bacterial hosts. Notably, the moderate purity of the host range within the order *Bifidobacteriales* may stem from the presence of broad-host-range (BHR) plasmids, which can be hosted by bacteria across different orders.

**Fig. 9.**
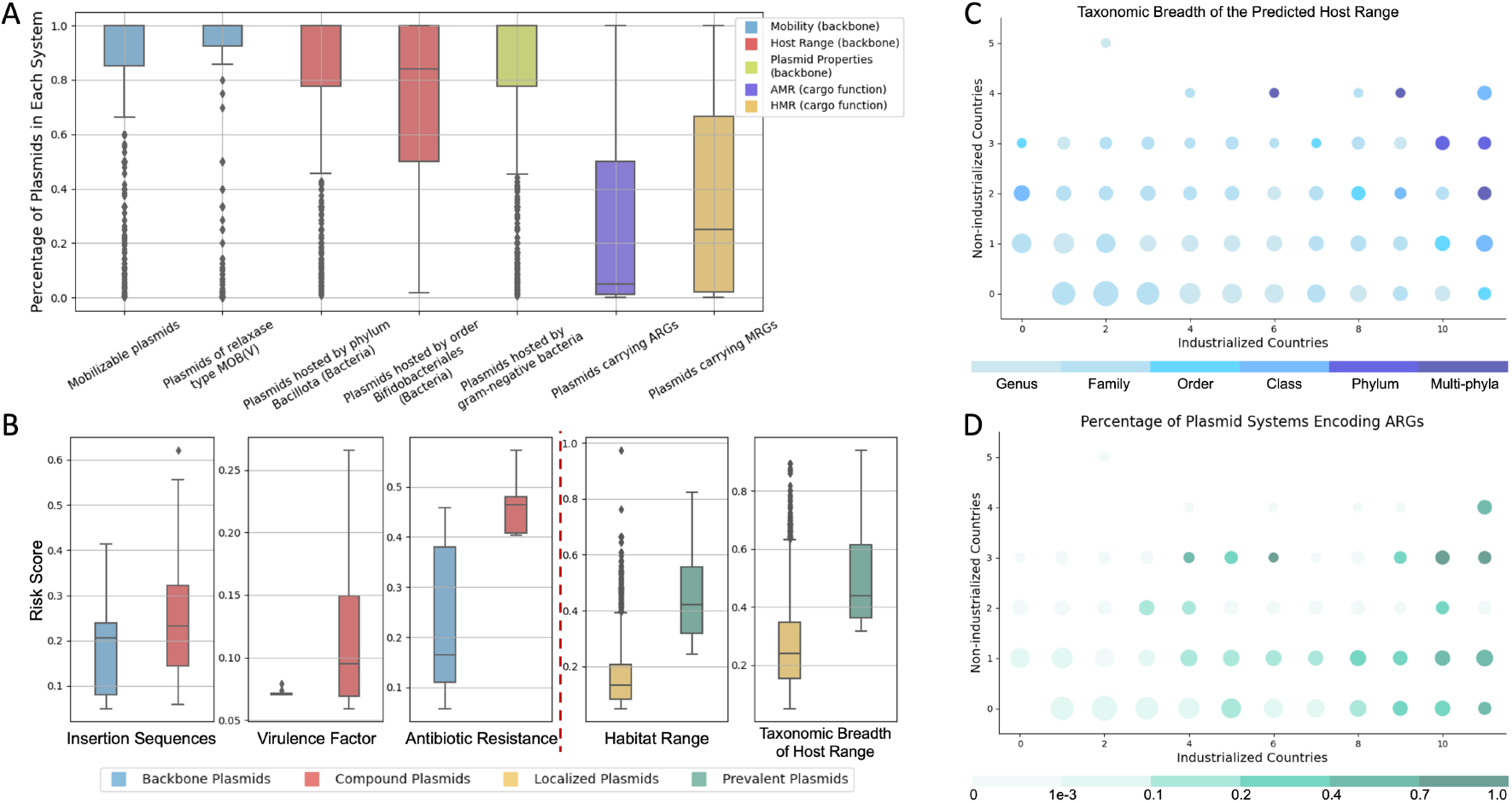
The results analyzed by PlasRAG across 1,169 PSs. (A) Distribution of ‘property-specific purity’ of seven representative properties across all PSs. Purity is calculated as the percentage of plasmids predicted by each property within a PS. Notably, PSs with a purity of 0 were excluded from the box plot, as this indicates that the property is irrelevant. The overall high percentage values suggest that the property is more closely associated with plasmid backbone functions. (B) Distribution comparison of different plasmid groups based on risk index across five sub-categories. It is reasonable to conclude that compound plasmids pose more risk than backbone plasmids in terms of their encoded ISs, VFs, and ARGs. Additionally, we observed that plasmids with broader dissemination tend to pose greater risks regarding their bacterial host range breadth and source habitat diversity. (C) The most common taxonomic level of host range among various PSs, grouped by their number of source countries (industrialized and non-industrialized). The circle size represents the number of PSs within each group, while the breadth of host range is defined by the most common taxonomic level in that group. (D) Percentage of PSs encoding ARGs within each lifestyle-based PS group. The display style is consistent with (C). (C) and (D) collectively suggest that plasmids with a broad host range and encoded ARGs are more likely to be present in a greater diversity of countries.

#### Insights gained from risk index comparisons

*Risk Index* is another crucial facet for mining insights about plasmids revealed from real human gut metagenomes. Therefore, we evaluated the distribution of the risk index among different plasmid groups classified by two schemes. One scheme categorizes plasmids as backbone or compound plasmids, while the other is prevalence-based, determined by the lifestyles of the source individuals. Specifically, we defined plasmids found in both non-industrialized and industrialized countries as prevalent plasmids, while those exclusive to one type of country were classified as localized plasmids. As shown in Fig. 9B, five groups of comparisons were observed across five *Risk Index* subcategories. First, the risk indexes evaluated based on the number of insertion sequences (ISs), ARGs, and VFs were higher for compound plasmids than for backbone plasmids, with clearer distinctions observed in ARGs and VFs. This is expected, as AMR and VFs are key accessory traits encoded on plasmids. On the other hand, while ISs are typically not considered part of plasmid backbones, they are conserved for specific cases of plasmid evolution [49]. Second, a clear distinction was observed in both risk indexes assessed based on host habitats and the breadth of bacterial host range between localized and prevalent plasmids. This suggests that plasmids present in a wider range of ecosystems and with a broader host range are more common or can be transferred to a greater diversity of bacteria, thereby becoming prevalent in various human societies.

#### Factors influencing plasmid ecology

Further analysis indicated that the distribution of plasmids among different source individuals is associated with two factors: host range and ARGs. As detailed in the ‘Methods’ section, PlasRAG can predict the taxonomic levels of the plasmid host range (e.g., the family level indicates that the plasmids can be hosted by bacteria of multiple genera within the family), reflecting its breadth. Thus, we employed PlasRAG to identify the most common taxonomic level of host range in each PS. We then presented the results in Fig. 9C, grouping the PSs by their number of source countries from non-industrialized and industrialized regions. A positive correlation was observed between the breadth of plasmid host range and the prevalence of plasmids across different countries. For instance, a broad host range encompassing multiple orders within a single phylum frequently occurred for PSs sourced from at least 10 countries. This *Host Range* analysis offered an additional perspective to explain the weak correlation between the distribution of plasmids and their bacterial hosts at the species level, as discussed by Yu et al. in [46]. Second, we evaluated the correlation between the prevalence of plasmids and the ARGs they encode. As illustrated in Fig. 9D, we displayed the percentage of PSs encoding ARGs for each source group, using the same style as in Fig. 9C. Specifically, a low percentage of less than 0.1 was observed for all PSs sourced from fewer than five countries. In contrast, a percentage of ARGs exceeding 0.2 frequently occurred in PSs found in more than eight countries. A country-specific analysis revealed that ARG-encoded plasmids originating from China and Spain are more abundant than those from Australia and Denmark. For instance, 72 out of 954 plasmids sourced from China (7.55%) encode ARGs resistant to 16 antibiotics, whereas only 6 out of 480 plasmids from Denmark encode ARGs resistant to 4 antibiotics. This finding aligns with the observation in [50] that the abundance of ARGs is significantly correlated with per capita antibiotic usage rates in each country. Therefore, it is reasonable to conclude that plasmids are one of the primary carriers of ARGs within microbial diversity. In summary, the PlasRAG results on the PS dataset indicated that plasmids with a broader host range and a greater number of encoded ARGs tend to spread more extensively, thereby having a more significant impact on environmental and public health.

## Discussion

In this study, we designed PlasRAG, a deep learning-based tool that realizes two efficient modules: plasmid property characterization and plasmid DNA retrieval across 10 important property facets, such as ecosystems and risk index. The core idea of PlasRAG is to treat property annotations as text and to design a multi-modal learning algorithm that aligns the plasmid sequence-text correlations between the two modalities. Leveraging the rich knowledge from textual properties provided by biomedical LMs, PlasRAG can merge divergent annotations from different reference databases while enhancing performance in sequence-text alignment. Furthermore, the sequence-to-text (S2T) characterization module is built upon the RAG pipeline. By augmenting a prompt with retrieved properties from the multi-modal IR model, the most relevant literature abstract, and the user query, the powerful LLM Llama-3 can generate more comprehensive and accurate responses. We conducted a series of experiments to validate the effectiveness of PlasRAG’s design. First, the benchmark experiments and embedding visualization demonstrated that the multi-modal architecture effectively learns the associations between plasmid sequences and property texts. Second, a detailed presentation of the conversational outputs highlights the usage and design rationale of PlasRAG. Finally, the multi-faceted characterization results on the novel PS dataset, sourced from diverse human gut metagenomes, provide several new ecological insights about plasmids.

Although we have shown the advantages of PlasRAG in several application scenarios, there is still potential for improvement. First, we exclusively extract sequence features from the CDSs in plasmids, as proteins commonly encode key genetic information closely associated with plasmid properties and functionality. However, non-coding regions may also contain valuable information. Due to the high scalability of the Perceiver Resampler’s cross-attention mechanism, an advanced design that introduces the long-context genomic foundation model, Evo 2 [51], to supplement non-coding features could enhance sequence-text alignment performance. Second, our work provides a framework primarily based on multi-modal learning and the RAG pipeline, which is not limited to plasmid analysis. Instead, our lightweight and scalable model architecture can also be extended to other biological entities, such as bacteria. Finally, as detailed in the Methods section, there are extensive plasmid-related literature resources (3,120 publications) available in NCBI RefSeq. Therefore, employing the powerful LLM agents to summarize high-quality and novel property labels, rather than relying on manual curation, may create significant opportunities for exploration.

