## Supplemental Data 1 for "PlasRAG: comprehensive plasmid characterization and retrieval through sequence-text alignment"

### Supplementary information for "PlasRAG: comprehensive plasmid characterization and retrieval through sequence-text alignment"

Yongxin Ji, Herui Liao, Jiaojiao Guan, Jiayu Shang, and Yanni Sun

Electrical Engineering Department, City University of Hong Kong, Kowloon, Hong Kong SAR

May 2025

#### 1 Details of Curated Textual Property Data

**Supplementary Table S1.** The containment relationship between the 10 property facets and the three databases, together with brief descriptions of the property contents. The 24 risk index properties labeled with a star can be categorized into four risk levels corresponding to six risk subcategories.

| Facet (# of properties) | Existence of annotations | | | Description of contents |
|---|---|---|---|---|
| | PIPdb | IMG/PR | PLSDB | |
| Antimicrobial resistance (124) | ✓ | ✓ | ✓ | Resistance mechanisms, antimicrobial agents, and antimicrobial resistance genes (ARGs) prioritized by WHO, along with antibiotic classes to which ARGs can confer resistance. |
| Bacterial host range (199) | ✓ | ✓ | ✓ | Host ranges represented by bacterial taxonomic labels standardized by the NCBI Taxonomy Database. |
| Ecological host (43) | × | × | ✓ | The taxonomic labels of host-associated ecosystems standardized by the NCBI Taxonomy Database. |
| Ecosystems (58) | ✓ | ✓ | ✓ | Ecosystems classified according to two schemes (PLSDB and IMG/M), along with pathogen habitats under the PIPdb scheme. |
| Fundamental property (15) | ✓ | ✓ | ✓ | Completeness, topology, data source, gram-staining reaction of bacterial hosts, and phage-plasmid identification. |
| Heavy metal resistance (231) | ✓ | × | × | Metal resistance genes (MRGs) and the metals to which they confer resistance. |
| Incompatibility group (59) | ✓ | × | ✓ | Incompatibility (Inc) groups according to three schemes: PlasmidFinder, Plasmid MLST, and MOB-typer. |
| Mobility (41) | × | ✓ | ✓ | Three plasmid mobility categories, relaxase types according to two schemes (CONJScan and MOB-typer), mating pair formation (MPF) types according to two schemes (CONJScan and MOB-typer), and T4SS ATPases along with Type IV coupling proteins (T4CPs). |
| Risk index (24*) | ✓ | × | × | Risk indexes assessed from six subcategories. |
| Virulence factor (205) | ✓ | × | × | Specific virulence factors (VFs) and 13 primary VF categories. |

**Supplementary Table S2.** Detailed list of our curated 10-faceted vocabulary. Due to space limitations, we present most textual properties using sentence templates highlighted in red, together with specific labels that can serve as candidates for the *label* in these templates.

| | | | | | | | |
|---|---|---|---|---|---|---|---|
| **Antimicrobial resistance - 1:** The plasmid encodes ARGs that confer resistance to the drug class **{label}.** | | | | | | | |
| aminocoumarin antibiotic | | aminoglycoside antibiotic | | diaminopyrimidine antibiotic | | disinfecting agents and antiseptics | |
| fluoroquinolone antibiotic | | fusidane antibiotic | | glycopeptide antibiotic | | lincosamide antibiotic | |
| macrolide antibiotic | | mupirocin-like antibiotic | | nitrofuran antibiotic | | nucleoside antibiotic | |
| oxazolidinone antibiotic | | peptide antibiotic | | phenicol antibiotic | | phosphonic acid antibiotic | |
| pleuromutilin antibiotic | | rifamycin antibiotic | | streptogramin antibiotic | | sulfonamide antibiotic | |
| sulfone antibiotic | | tetracycline antibiotic | | carbapenem | cephalosporin | cephamycin | monobactam |
| penam | penem | | | | | | |
| **Antimicrobial resistance - 2:** The plasmid encodes the WHO-prioritized ARG **{label}**. | | | | | | | |
| ErmB | IMP-1 | IMP-4 | MecA | NDM-1 | OXA-10 | OXA-181 | OXA-23 |
| OXA-24 | OXA-48 | OXA-58 | QacE | QnrA1 | QnrB19 | QnrS1 | SHV-1 |
| TEM-1 | Tet(M) | VIM-1 | VIM-2 | VIM-4 | VanA | VanB | |
| **Antimicrobial resistance - 3:** The plasmid encodes the antibiotic resistance mechanism **{label}**. | | | | | | | |
| ABC Transporter | | Acetyltransferase | | Aminoglycoside Modifying Enzyme | | Antibiotic Inactivation | |
| Chloramphenicol Resistance | | Class A Beta-Lactamase | | Class B Beta-Lactamase | | Class C Beta-Lactamase | |
| Class D Beta-Lactamase | | Gene Modulating Resistance | | Gylcopeptide Resistance | | MFS Transporter | |
| Nucleotidyltransferase | | Phosphotransferase | | Quinolone Resistance | | RND Antibiotic Efflux | |
| Target Protection | | Tetracycline Inactivation | | Tetracycline MFS Efflux | | Tetracycline Ribosomal Protection | |
| rRNA Methyltransferase | | Other Efflux | | | | | |
| **Antimicrobial resistance - 4:** The plasmid can confer antimicrobial resistance to the agent **{label}**. | | | | | | | |
| chloramphenicol | | quaternary ammonium | | aminoglycoside | | streptogramin b | |
| streptothricin | beta-lactam | quinolone | lincosamide | nickel | clindamycin | apramycin | tylosin |
| vancomycin | copper | florfenicol | gentamicin | mupirocin | glycopeptide | tiamulin | streptomycin |
| tobramycin | amikacin | tellurium | arsenic | oxazolidinone | macrolide | silver | arsenate |
| phenicol | erythromycin | bleomycin | rifamycin | virginiamycin | kanamycin | hygromycin | organomercury |
| streptogramin | telithromycin | sulfonamide | cadmium | azithromycin | linezolid | tetracycline | spiramycin |
| arsenite | colistin | mercury | fosfomycin | tigecycline | pleuromutilin | trimethoprim | |
| **Bacterial host range - 1:** The plasmid is hosted by bacteria in the phylum **{label}**. | | | | | | | |
| Actinomycetota | | Campylobacterota | | Cyanobacteriota | | Fusobacteriota | |
| Methanobacteriota | | Mycoplasmatota | | Pseudomonadota | | Thermodesulfobacteriota | |
| Verrucomicrobiota | | Bacillota | Bacteroidota | Chlamydiota | Deinococcota | Spirochaetota | |
| **Bacterial host range - 2:** The plasmid is hosted by bacteria in the class **{label}**. | | | | | | | |
| Alphaproteobacteria | | Betaproteobacteria | | Epsilonproteobacteria | | Erysipelotrichia | |
| Flavobacteriia | | Gammaproteobacteria | | Actinomycetes | Bacilli | Bacteroidia | Clostridia |
| Cyanophyceae | Cytophagia | Deinococci | Fusobacteriia | Halobacteria | Spirochaetia | | |
| **Bacterial host range - 3:** The plasmid is hosted by bacteria in the order **{label}**. | | | | | | | |
| Alteromonadales | | Bifidobacteriales | | Burkholderiales | | Campylobacterales | |
| Enterobacterales | | Erysipelotrichales | | Flavobacteriales | | Fusobacteriales | |
| Halobacteriales | | Hyphomicrobiales | | Kitasatosporales | | Lachnospirales | |
| Lactobacillales | | Lysobacterales | | Mycobacteriales | | Pasteurellales | |
| Peptostreptococcales | | Pseudomonadales | | Rhodobacterales | | Rhodospirillales | |
| Sphingomonadales | | Spirochaetales | | Neisseriales | Nostocales | Thiotrichales | Vibrionales |
| Aeromonadales | Bacillales | Bacteroidales | Cytophagales | Deinococcales | Eubacteriales | Legionellales | Leptospirales |
| Micrococcales | Moraxellales | | | | | | |
| **Bacterial host range - 4:** The plasmid is hosted by bacteria in the family **{label}**. | | | | | | | |
| Acetobacteraceae | | Aeromonadaceae | | Alcaligenaceae | | Bacteroidaceae | |
| Bifidobacteriaceae | | Burkholderiaceae | | Campylobacteraceae | | Clostridiaceae | |
| Comamonadaceae | | Corynebacteriaceae | | Enterobacteriaceae | | Enterococcaceae | |
| Fusobacteriaceae | | Haloferacaceae | | Helicobacteraceae | | Lachnospiraceae | |
| Lactobacillaceae | | Legionellaceae | | Leptospiraceae | | Lysobacteraceae | |
| Methylobacteriaceae | | Microbacteriaceae | | Micrococcaceae | | Morganellaceae | |
| Mycobacteriaceae | | Nitrobacteraceae | | Oscillospiraceae | | Paenibacillaceae | |
| Pasteurellaceae | | Pectobacteriaceae | | Peptostreptococcaceae | | Phyllobacteriaceae | |
| Piscirickettsiaceae | | Prevotellaceae | | Pseudomonadaceae | | Roseobacteraceae | |
| Sphingomonadaceae | | Staphylococcaceae | | Streptococcaceae | | Streptomycetaceae | |
| Bacillaceae | Borreliaceae | Brucellaceae | Erwiniaceae | Listeriaceae | Moraxellaceae | Neisseriaceae | Nocardiaceae |
| Nostocaceae | Paracoccaceae | Rhizobiaceae | Vibrionaceae | Yersiniaceae | | | |
| **Bacterial host range - 5:** The plasmid is hosted by bacteria in the genus **{label}**. | | | | | | | |
| Bradyrhizobium | | Clostridioides | | Corynebacterium | | Faecalibacterium | |
| Komagataeibacter | | Lacticaseibacillus | | Lactiplantibacillus | | Ligilactobacillus | |

| | | | | | | | |
|---|---|---|---|---|---|---|---|
| Methylobacterium | | Mycobacteroides | | Mycolicibacterium | | Novosphingobium | |
| Paraburkholderia | | Photobacterium | | Piscirickettsia | | Staphylococcus | |
| Stenotrophomonas | | Streptococcus | Streptomyces | Tritonibacter | Vibrio | Xanthomonas | Yersinia |
| Acetobacter | Acinetobacter | Aeromonas | Agrobacterium | Bacillus | Bacteroides | Bordetella | Borrelia |
| Borreliella | Brucella | Burkholderia | Campylobacter | Citrobacter | Clostridium | Cronobacter | Ensifer |
| Enterobacter | Enterococcus | Erwinia | Escherichia | Fusobacterium | Helicobacter | Klebsiella | Lactobacillus |
| Lactococcus | Legionella | Leptospira | Leuconostoc | Listeria | Mesorhizobium | Micrococcus | Moraxella |
| Morganella | Mycobacterium | Neisseria | Nocardia | Nostoc | Pantoea | Paracoccus | Phaeobacter |
| Phocaeicola | Prescottella | Prevotella | Priestia | Proteus | Providencia | Pseudomonas | Ralstonia |
| Rhizobium | Rhodococcus | Salmonella | Segatella | Serratia | Shigella | Sinorhizobium | Sphingobium |
| Sphingomonas | | | | | | | |
| **Ecological host - 1:** The plasmid can be found in ecosystems associated with hosts of the phylum **{label}**. | | | | | | | |
| Arthropoda | Bacillariophyta | Chordata | Streptophyta | | | | |
| **Ecological host - 2:** The plasmid can be found in ecosystems associated with hosts of the class **{label}**. | | | | | | | |
| Actinopteri | Arachnida | Aves | Insecta | Magnoliopsida | Malacostraca | Mammalia | |
| **Ecological host - 3:** The plasmid can be found in ecosystems associated with hosts of the order **{label}**. | | | | | | | |
| Anseriformes | Artiodactyla | Carnivora | Decapoda | Fabales | Galliformes | Hymenoptera | Ixodida |
| Poales | Primates | Rodentia | Rosales | Salmoniformes | Solanales | | |
| **Ecological host - 4:** The plasmid can be found in ecosystems associated with hosts of the family **{label}**. | | | | | | | |
| Bovidae | Canidae | Fabaceae | Hominidae | Penaeidae | Phasianidae | Poaceae | Salmonidae |
| Solanaceae | Suidae | | | | | | |
| **Ecological host - 5:** The plasmid can be found in ecosystems associated with hosts of the genus **{label}**. | | | | | | | |
| Bos | Canis | Gallus | Homo | Sus | | | |
| **Ecological host - 6:** The plasmid can be found in ecosystems associated with hosts of the **{label}**. | | | | | | | |
| species Homo sapiens | | superkingdom Bacteria | | superkingdom Eukaryota | | | |
| **Ecosystem range - 1:** The plasmid can be found in pathogens associated with the **{label}** habitat. | | | | | | | |
| Arthropod | Birds | Environment | Food | Human | Mammal | Other | Rodent |
| **Ecosystem range - 2:** The plasmid can be found in the **{label}** ecosystem, according to the PLSDB scheme. | | | | | | | |
| anthropogenic | aquatic | cell culture | estuary | food | host associated | location | marine |
| saline | sea | spring | | | | | |
| **Ecosystem range - 3:** The plasmid can be found in the **{label}** ecosystem, according to the IMG/M scheme. | | | | | | | |
| simulated communities (contig mixture) | | | | simulated communities (microbial mixture) | | | |
| arthropoda: insects | | digestive system | | host-associated | | hydrothermal vents | |
| large intestine | | mammals: human | | non-marine saline and alkaline | | reproductive system | |
| rock-dwelling (endoliths) | | ssf (solid state fermentation) | | terrestrial | tundra | vagina | stomach |
| algae | anaerobic | annelida | aquatic | engineered | environmental | floodplain | freshwater |
| gut | larva | mammals | marine | modeled | oral cavity | palsa | peat |
| peat moss | plants | porifera | roots | soil | solar panel | sponge | |
| **Fundamental property - 1:** The plasmid is sourced from **{label}**. | | | | | | | |
| Metagenome-assembled genomes (MAGs) | | | | Single amplified genomes (SAGs) | | | |
| Isolates | | Metagenomes | | Metatranscriptomes | | | |
| **Fundamental property - 2:** Completeness, topology, gram-staining reaction of bacterial hosts, and phage-plasmid identification. | | | | | | | |
| Complete plasmid. | | Fragmented plasmid. | | Circular plasmid. | | Linear plasmid. | |
| The plasmid contains concatemer structures. | | | | The plasmid contains direct terminal repeat structures. | | | |
| The plasmid contains inverted terminal repeat structures. | | | | Phage-plasmid. | | | |
| The plasmid is hosted by Gram-negative bacteria. | | | | The plasmid is hosted by Gram-positive bacteria. | | | |
| **Heavy metal resistance - 1:** The plasmid encodes genes that confer resistance to **{label}**. | | | | | | | |
| Aluminium (Al) | Antimony (Sb) | Arsenic (As) | Bismuth (Bi) | Cadmium (Cd) | Chromium (Cr) | Cobalt (Co) | Copper (Cu) |
| Gallium (Ga) | Gold (Au) | Iron (Fe) | Lead (Pb) | Magnesium (Mg) | Manganese (Mn) | Mercury (Hg) | Molybdenum (Mo) |
| Nickel (Ni) | Selenium (Se) | Silver (Ag) | Tellurium (Te) | Tungsten (W) | Vanadium (V) | Zinc (Zn) | |
| **Heavy metal resistance - 2:** The plasmid encodes the metal resistance gene **{label}**. | | | | | | | |
| G2alt | acn | acr3 | actP | actR | aioA (aoxB) | aioE | aioR (aoxR) |
| arsA | arsB | arsC | arsD | arsH | arsM | arsP | arsR |
| arsT | baeR | baeS | bhsA (comC, ycfR) | cadC | cadD | cadX | chrA |
| chrA1 | chrB | chrB1 | chrF | chrR | cmtR | comR (ycfQ) | cop-unnamed |
| copA | copB | copC | copD | copF | copG | copM | copR |
| copS | copY (tcrY) | copZ | corA | corC | corR | corR (coaR) | corS |
| corT (coaT) | crdR | csoR | ctpV | cueA | cueP | cusA (ybdE) | cusB |
| cusC (ylcB) | cusF (cusX) | cusR (ylcA) | cusS | cutA | czcA | czcB | czcC |
| czcD | czcR | czcS | dmeF | dmeR | dpsA | dsbA | dsbC |
| fbpB | fbpC | fecD | fecE | fetA (ybbL) | fetB (ybbM) | fieF (yiip) | fptA |
| fpvA | gesA | gesB | gesC | glpF | golS | golT | hmrR |
| hoxN | irlR | irlS | klaB (kilB, telA) | klaC (telB) | kmtR | mco | mdtA |
| merA | merB | merC | merD | merE | merF | merG | merP |

| | | | | | | | |
|---|---|---|---|---|---|---|---|
| merR | merR1 | merR2 | merT | mgtA | mntH (yfeP) | modA | modB |
| modC | mreA | ncrA | ncrB | ncrC | ncrY | nikA | nikB |
| nikC | nikD | nikE | nirD | nixA | nrsD (nreB) | nrsS | pbrA |
| pbrB (pbrC) | pbrR | pbrT | pcoA | pcoB | pcoC | pcoD | pcoE |
| pcoR | pcoS | pfr | pgpA (ltpgpA) | pitA | pmrA | pmrB | pmrC |
| pmrG | pstA | pstB | pstC | pstS | rcnA (yohM) | rcnB (yohN) | rcnR (yohL) |
| recG | ricR | robA | ruvB | silA | silB | silC | silE |
| silF | silP | silR | silS | sitA | sitB | sitC | sitD |
| smtB (ziaR) | soxS | tcrA | tcrB | tcrY | tcrZ | tehA | tehB |
| terA | terB | terC | terD | terE | terW | terZ | troB |
| tunR | tupC | wtpC | ybtP | ybtQ | yfeA | yfeB | yfeD |
| yfmP | ygiW | yhcN | yieF | yjaA | yodD | yqjH | zevB |
| ziaR | zinT (yodA) | zitB (ybgR) | zntR (yhdM) | znuC (yebM) | zraR (hydH) | zraS (hydG) | zur (yjbK) |
| **Incompatibility group - 1:** The plasmid belongs to the **{label}** incompatibility group, according to the PlasmidFinder scheme. | | | | | | | |
| IncA/C2 | IncB/O/K/Z | IncFIA | IncFIB | IncFIC | IncFII | IncHI1A | IncHI1B |
| IncHI2 | IncHI2A | IncI | IncI1 | IncI2 | IncL/M | IncN | IncN2 |
| IncP1 | IncP6 | IncQ1 | IncR | IncU | IncX1 | IncX2 | IncX3 |
| IncX4 | IncX5 | IncY | | | | | |
| **Incompatibility group - 2:** The plasmid belongs to the **{label}** incompatibility group, according to the Plasmid MLST scheme. | | | | | | | |
| IncA/C | IncF | IncHI1 | IncHI2 | IncI1 | IncN | | |
| **Incompatibility group - 3:** The plasmid belongs to the **{label}** incompatibility group, according to the MOB-typer scheme. | | | | | | | |
| Inc11 | Inc13 | Inc18 | IncC | IncFIA | IncFIB | IncFIC | IncFII |
| IncHI1A | IncHI1B | IncHI2A | IncI-gamma/K1 | IncI1 | IncI1/B/O | IncI2 | IncK2/Z |
| IncL/M | IncN | IncP | IncQ1 | IncR | IncU | IncX1 | IncX3 |
| IncX4 | IncY | | | | | | |
| **Mobility - 1:** Three plasmid mobility categories. | | | | | | | |
| Conjugative plasmid. | | Mobilizable plasmid. | | Non-mobilizable plasmid. | | | |
| **Mobility - 2:** The plasmid is transferable and belongs to mobilization type **{label}**, according to the CONJScan scheme. | | | | | | | |
| MOBB | MOBC | MOBF | MOBH | MOBP1 | MOBP2 | MOBP3 | MOBQ |
| MOBT | MOBV | | | | | | |
| **Mobility - 3:** The plasmid is transferable and belongs to mobilization type **{label}**, according to the MOB-typer scheme. | | | | | | | |
| MOBB | MOBC | MOBF | MOBH | MOBM | MOBP | MOBQ | MOBT |
| MOBV | MOB_Unknown | | | | | | |
| **Mobility - 4:** The plasmid encodes a complete conjugation system and belongs to the mating-pair-formation type **{label}**, according to the CONJScan scheme. | | | | | | | |
| MPF_B | MPF_C | MPF_F | MPF_FA | MPF_FATA | MPF_G | MPF_I | MPF_T |
| **Mobility - 5:** The plasmid encodes a complete conjugation system and belongs to the mating-pair-formation type **{label}**, according to the MOB-typer scheme. | | | | | | | |
| MPF_F | MPF_I | MPF_T | MPF_Unknown | | | | |
| **Mobility - 6:** The plasmid encodes the **{label}** within the T4SS conjugation system. | | | | | | | |
| T4SS ATPase F_traU | | T4SS ATPase I_traU | | T4SS ATPase virb4 | | Type IV coupling protein t4cp1 | |
| Type IV coupling protein t4cp2 | | Type IV coupling protein tcpA | | | | | |
| **Risk index:** The risk index of the plasmid assessed based the **{label}**. | | | | | | | |
| Number of insertion sequences harbored by the plasmid | | | | Distribution of the plasmid across different habitats | | | |
| Number of virulence factor genes harbored by the plasmid | | | | Number of ARGs harbored by the plasmid | | | |
| Number of ARGs from WHO Priority List harbored by the plasmid | | | | Taxonomic breadth of the plasmid host range | | | |
| **Virulence factor - 1:** The plasmid carries virulence factors that belong to the **{label}** category. | | | | | | | |
| Antimicrobial activity/Competitive advantage | | | | Nutritional/Metabolic factor | | | |
| Effector delivery system | | | | Immune modulation | | | |
| Adherence | | Biofilm | | Exoenzyme | | Exotoxin | |
| Invasion | | Motility | | Other | | Regulation | |
| Stress survival | | | | | | | |
| **Virulence factor - 2:** The plasmid carries the virulence factor **{label}**. | | | | | | | |
| Bee (biofilm enhancer in enterococci) | | | | Contact-dependent inhibition CDI system | | | |
| Enterobactin synthesis and transport | | | | Locus for diffuse adherence (lda), afimbrial adhesin | | | |
| Lvh (Legionella vir homologs) type IVA secretion system | | | | PilA-type pili (PGS1, pilin gene clusters 1) | | | |
| Vacuolating autotransporter gene Vat | | | | Yersiniabactin | | Yersiniabactin siderophore | |
| <alpha>-Hemolysin | | AAI/SCI-II T6SS | | AatA, AIDA-I type | | Adhesive fimbriae | |
| Afimbrial adhesin AFA-I | | Agglutinin receptor | | Allantion utilization | | Antigen 43, AIDA-I type | |
| Bsa T3SS secreted effectors | | CS31A capsule-like antigen | | Cah, AIDA-I type | | Clumping factor | |
| Direct heme uptake system | | Dot/Icm T4SS secreted effectors | | EhaA, AIDA-I type | | EhaB, AIDA-I type | |
| Enterotoxin SenB/TieB | | Ferrous iron transport | | GacS/GacA two-component system | | Hbp (hemoglobin-binding protease) | |
| Heat-labile toxin (LT) | | Heat-stable toxin (ST) | | Hyaluronate lyase | | Hyaluronic acid (HA) capsule | |
| Insecticidal crystalline toxins | | Intercellular adhesion proteins | | Iron-regulated element | | Iron/manganese transport | |
| Peritrichous flagella | | PilB-type pili (PGS3) | | Quorum-sensing | | SCI (Salmonella centrisome island) | |
| Salmochelin siderophore | | Serine protease splABCDEF | | Staphyloferrin A | | T3SS1 secreted effectors | |

| | | | | | | | |
|---|---|---|---|---|---|---|---|
| T3SS2 secreted effectors | | TTSS (SPI-1 encode) | | TTSS (SPI-2 encode) | | TTSS secreted effectors | |
| TTSS-1 secreted effectors | | TTSS-2 secreted effectors | | Thermonuclease nuc | | Type 1 fimbriae | |
| Type 3 fimbriae | | Type I fimbriae | | Type IV secretion system | | Type VII secretion system | |
| AAFs | ACE T6SS | AS | Aerobactin | Afa/Dr family | Agf | AggR | Anthrax toxin |
| ApeE | AtxA | Aureolysin | Autolysin | BFP | Bcf | BimA | Bsa T3SS |
| BslA | C3610 | CDT | CNF-1 | Capsule | ClyA | Col10 | Col5 |
| Colibactin | Colicin B | Colicin E1 | Colicin Ia | Colicin Ib | Colicin K | Colicin N | Colicin S4 |
| Colicin U | Colicin Y | Csu fimbriae | Curli fibers | Cytolysin | Dispersin | Dr adhesins | ECP |
| EHS | ESX-1 | ESX-3 | ESX-5 | ETT2 | EaeH | Ebp pili | Efa-1/LifA |
| Emp | Ent | Enterobactin | EpeA | EspC | EspI, SPATE | EspP | Etp |
| EtpA | F1C fimbriae | F9 fimbriae | Flagella | FnBPs | Gsp | HBL | HemO cluster |
| Hemolysin | Ibes | IcsA (VirG) | IcsP (SopA) | IgA1 protease | Intimin | K1 capsule | LPS |
| Ler | Lpf | LukED | MgtBC | Mig-14 | Mig-5 | MisL | MsbB2 |
| OmpD | P fimbriae | Paa | PagN | PagR-XO1 | Pef | Pet | Pic |
| Pix pilus | Pyoverdine | RatB | Rck | RcsAB | RmpA | RpoS | S fimbriae |
| SCI-I T6SS | SDr | SE | Saf | Sal | Salmochelin | Sat | Sfp fimbriae |
| ShET2 | ShdA | SigA | SinH | SpvB | Stb | StcE | Std |
| Stg fimbriae | T2SS | T3SS | T3SS1 | T3SS2 | T6SS | T6SS-1 | T6SS-II |
| T6SS-III | TFS3a | TFS3b | TTS1 | TTS2 | TTSS | Tcf | Tia/Hek |
| ToxB | TraJ | Tsh | Type IV pili | VirF | VirK | Ybt | |

#### 2 Computational Cost Evaluation Results

**Supplementary Table S3.** Required computational resources, including running time and maximum GPU memory, for both the training and prediction phases of the PlasRAG model. The evaluation focuses on the *Host Range* dataset detailed in Table 1 (main text), which typically incurs the highest computational costs. The PlasRAG model is lightweight, allowing us to utilize a large batch size of 2,048 for model training while using up to 64.6 GB of GPU memory. As expected, GPU memory usage increases linearly with batch size, while running time shows little variation. In particular, PlasRAG requires minimal time for predictions, enabling efficient usage for users.

| | Training Phase | | Prediction Phase | |
|---|---|---|---|---|
| Batch Size | Time | Maximum GPU Memory | Time | Maximum GPU Memory |
| 128 | 2h20min | 4.6GB | 40s | 3.0GB |
| 256 | 2h14min | 8.6GB | 39s | 5.3GB |
| 512 | 2h11min | 16.6GB | 39s | 10.1GB |
| 1024 | 2h11min | 32.6GB | 38s | 20.0GB |
| 2048 | 2h13min | 64.6GB | 38s | 38.6GB |

#### 3 Template for Augmenting Prompts: Integrating Properties, Literature, and User Queries

The following is the prompt template for augmented LLM generation: The black text represents fixed elements of the template, while the red text denotes variables that can be replaced with information retrieved for the query plasmids. The notation ***text*** indicates bold formatting in the Markdown prompt, serving to represent a subtitle.

Prompt Template for Augmented LLM Generation

You are an expert in plasmid analysis (bioinformatics) tasked with providing insights based on genetic information.

**Context**: We have analyzed a user-provided plasmid sequence and generated predictions regarding its properties from multiple facets. Additionally, we have identified the most relevant reference plasmid together with its associated literature abstract to support your answer.

**User Question**: Is the plasmid a multidrug-resistance plasmid? If so, please list all antibiotics to which it confers resistance.

**Predicted Properties**:
- **AMR**: the plasmid encodes ARGs that confer resistance to Beta-lactam (Cephalosporin, Monobactam, Penam), Bleomycin, Diaminopyrimidine, Fluoroquinolone (Quinolone), Nitrofuran, Phenicol, and Tetracycline (Glycylcycline). The associated resistance mechanisms include Class A beta-lactamase and RND antibiotic efflux. Additionally, the plasmid encodes the ARG TEM-1, listed in the WHO priority list;
- **Virulence Factor**: the plasmid carries the virulence factor Type 3 fimbriae, which belongs to the Biofilm category;
- **Metal Resistance**: the plasmid does not encode any metal resistance genes;
- **Host Range**: the plasmid is hosted by bacteria in the Enterobacteriaceae family;
- **Ecosystem**: the plasmid can be found in engineered ecosystems, ecosystems associated with the bird habitat, human habitat, and hosts of the genus Sus;
- **Mobility**: the plasmid is a conjugative plasmid, which encodes a complete conjugation system and belongs to the mating-pair-formation type MPF_T. It belongs to mobilization type MOBP1. Additionally, it encodes the T4SS ATPase virB4 and the Type IV coupling protein T4CP2 within the T4SS conjugation system;
- **Incompatibility Group**: the plasmid belongs to the incompatibility group IncX1;
- **Risk Index**: the plasmid has a combined Minimal risk level, categorized as follows: 1) Low risk based on its distribution across habitats, 2) Low risk based on its host range breadth, 3) Minimal risk based on virulence factor genes, 4) Minimal risk based on ARGs from WHO priority list, 5) Low risk based all encoded ARGs, and 6) Low risk based on insertion sequences;
- **Basic Properties**: the plasmid is a circular, complete plasmid sourced from an isolate, and hosted by Gram-negative bacteria.

**Retrieved Reference Plasmid**: pOLA52

**Literature Abstract**: The large conjugative multidrug resistance (MDR) plasmid pOLA52 was sequenced and annotated. The plasmid encodes two phenotypes normally associated with the chromosomes of opportunistic pathogens, namely MDR via a resistance-nodulation-division (RND)-type efflux-pump (oqxAB), and the formation of type 3 fimbriae (mrkABCDF). The plasmid was found to be 51,602 bp long with 68 putative genes. About half of the plasmid constituted a conserved IncX1-type backbone with predicted regions for conjugation, replication and partitioning, as well as a toxin/antitoxin (TA) plasmid addiction system. The plasmid was also classified as IncX1 with incompatibility testing. The conjugal transfer and plasmid maintenance regions of pOLA52 therefore seem to represent IncX1 orthologues of the well-characterized IncX2 plasmid R6K. Sequence homology searches in GenBank also suggested a considerably higher prevalence of IncX1 group plasmids than IncX2. The 21 kb 'genetic load' region of pOLA52 was shown to consist of a mosaic, among other things a fragmented Tn3 transposon encoding ampicillin resistance. Most notably the oqxAB and mrkABCDF cassettes were contained within two composite transposons (Tn6010 and Tn6011) that seemed to originate from Klebsiella pneumoniae, thus demonstrating the capability of IncX1 plasmids of facilitating lateral transfer of gene cassettes between different Enterobacteriaceae.

**Instructions**:
1. Ensure that your answer is clear and directly addresses the user's question, referencing the properties and literature as necessary when applicable.
2. Ensure that your answer strictly utilizes the details given and avoids any generation of additional information or hallucinations.
3. Avoid including simple information such as sequence length, gene count, or other basic metrics.
4. When analyzing the literature, summarize key information about the reference plasmid pOLA52 based on the abstract. Focus on specific functions, resistance characteristics, clinical significance, and mechanisms of action.

### 4 Instances Illustrating PlasRAG's Two Sub-modules

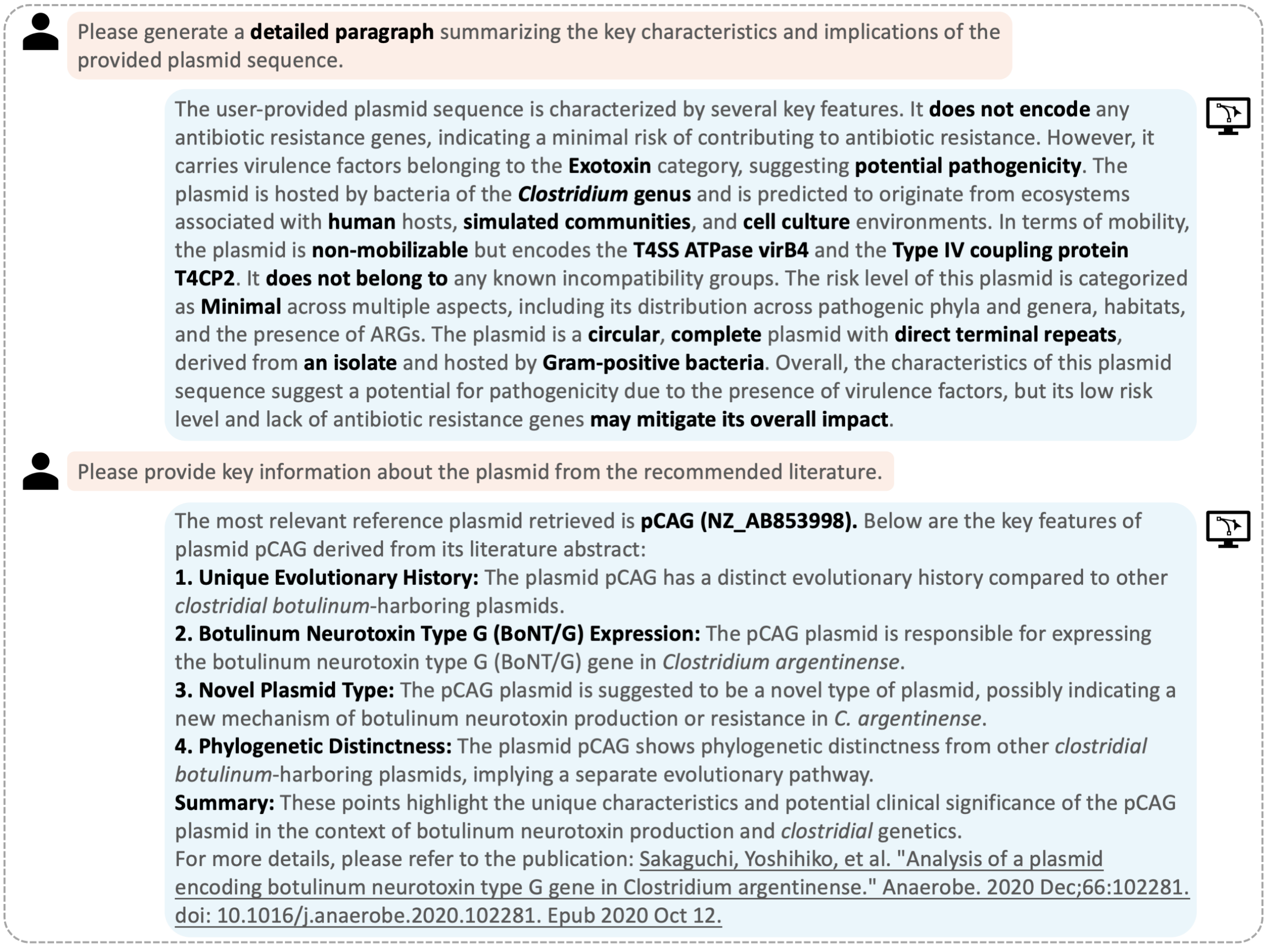


**Supplementary Figure S1.** LLM generation instances for characterizing the plasmid *IMGPR_plasmid_2772190775_000001* identified from the *Clostridium argentinense* 89G isolate.

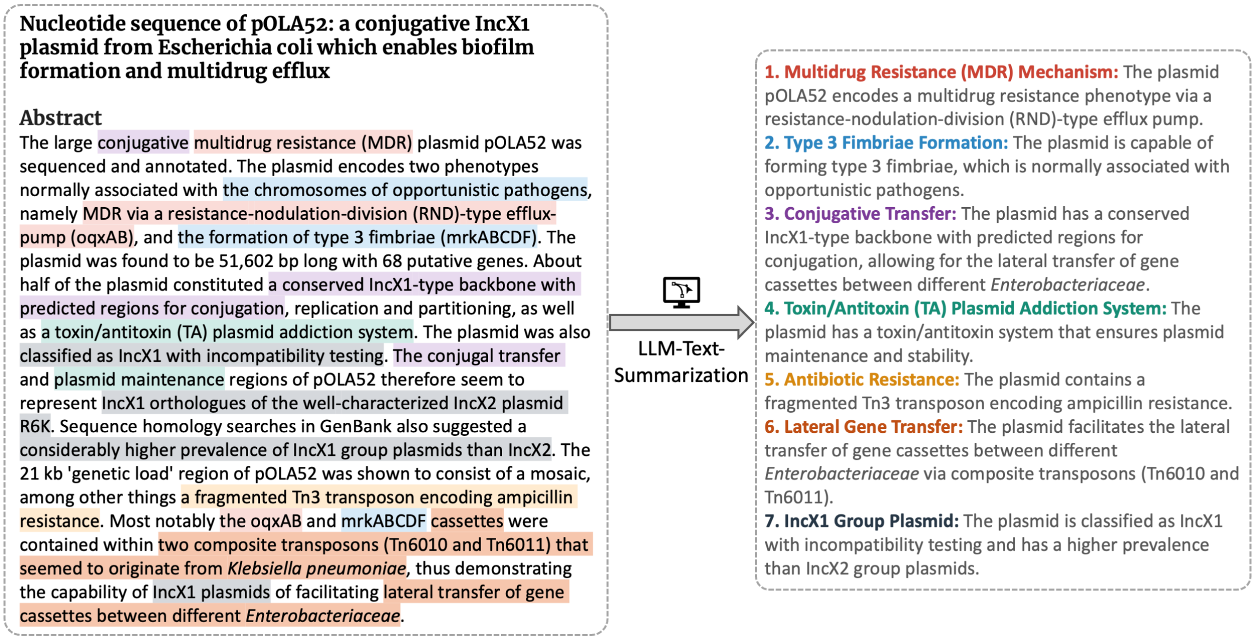


**Supplementary Figure S2.** Comparison between the literature abstract and the key knowledge list summarized by the LLM. Each of the seven points in the list is represented in specific colors: red, blue, purple, green, yellow, orange, and black. The sources for each point's knowledge are manually annotated with the corresponding colors.

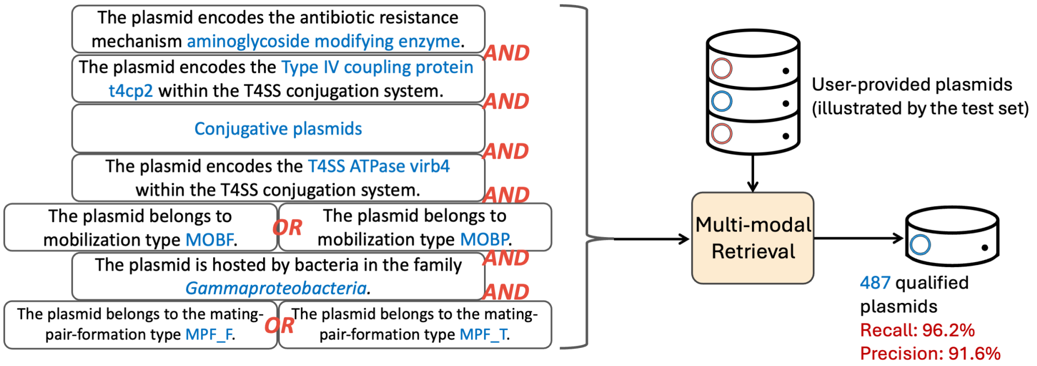


**Supplementary Figure S3.** An example of retrieving plasmids from a user-provided candidate set. The query condition includes nine selected properties combined with logical operators. The performance was evaluated by using the test set (100k plasmids) as the user-provided candidate set, resulting in a recall of 96.2% and precision of 91.6%.

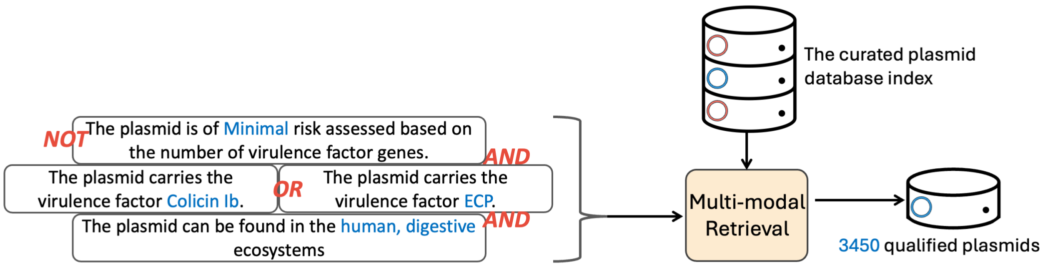


**Supplementary Figure S4.** An example of plasmid retrieval when no candidate set is provided by users. In this case, eligible plasmids are selected from our default indexed database.
